# Accurate prediction of functional states of *cis*-regulatory modules reveals the common epigenetic rules in humans and mice

**DOI:** 10.1101/2021.07.15.452574

**Authors:** Pengyu Ni, Joshua Moe, Zhengchang Su

## Abstract

We proposed a two-step approach for predicting active *cis*-regulatory modules (CRMs) in a cell/tissue type. We first predict a map of CRM loci in the genome using all available transcription factor binding data in the organism, and then predict functional states of all the putative CRMs in any cell/tissue type using few epigenetic marks. We have recently developed a pipeline dePCRM2 for the first step, and now presented machine-learning methods for the second step. Our approach substantially outperforms existing methods. Our results suggest common epigenetic rules for defining functional states of CRMs in various cell/tissue types in humans and mice.

## Backgrounds

Since the completion of the Human Genome Project 20 years ago [1, 2], we have largely categorized coding sequences in the genome [3] and gained a good understanding of their functions in various human cell/tissue types. In contrast, although *cis*-regulatory sequences can be as important as coding sequences in specifying human traits [4–6], our understanding of them falls largely behind due to more technical difficulties in categorizing them in the genome and in characterizing their functional states and target genes in various human cell/tissue types [7, 8]. *Cis*-regulatory sequences are also known as *cis*-regulatory modules (CRMs), as they often function independently of their locations and orientations relative to their target genes [9]. A CRM is generally made of a cluster of closely located transcription factor (TF) binding sites (TFBSs) in a genome region, to which cognate TFs can bind [10]. A CRM regulates gene transcription through interacting with the promoter of its target gene in a cell/tissue type where the constituent TFBSs in the CRM is accessible and the cognate TFs are expressed and available to bind [10, 11]. Thus, like a gene’s functional state (transcribed or untranscribed), a CRM’s functional state (TF-binding or non-TF-binding) is highly cell/tissue type specific.

Although traditional experimental methods for characterizing CRMs are highly accurate [12, 13], they are laborious and time consuming. Hence, various forms of massively parallel reporter assays (MPRA) have been developed [14]. In particular, self-transcribing assay of regulatory regions sequencing (STARR-seq) clones randomly sheared *D. melanogaster* genomic sequences between a minimal-promoter-driven GFP open reading frame and a downstream polyA sequence [15]. If a sequence is an active enhancer, this results in transcription of the enhancer sequence, allowing to assess more and relatively longer candidate sequences than earlier MPRA that used short (∼200bp) synthetic sequences [16]. Variants of STARR-seq have been developed to accommodate to large mammalian genomes, such as whole genome STARR-seq (WHG-STARR-seq) [17] and ATAC (assay for transposase-accessible chromatin) enrichment coupled with STARR-seq (ATAC-STARR-seq) [18]. However, since all forms of STARR-seq methods are based on episomal expression vectors, the results may not reflect the native chromosomal contexts [15–17, 19]. Moreover, sequences that can be assessed by STARR-seq are still much shorter (∼500bp) than the mean length (∼2,049bp) of known human enhancers in the VISTA database [13], they therefore suffer high false discovery rates (FDRs) as well as high false negative rates [15–22].

Since active (with TF-binding) and non-active (without TF-binding) CRMs in a cell/tissue type have distinct epigenetic marks [23–31], many machine-learning methods have been proposed to simultaneously predict CRM loci and their functional states in a given cell/tissue type based on genome segments’ chromatin accessibility (CA) as measured by DNase I hypersensitive sites sequencing (DNase-seq) [32] or assay for transposase accessible chromatin using sequencing (ATAC-seq) [33], histone modifications as measured by chromatin immunoprecipitation sequencing (ChIP-seq) [34] and cytosine methylation in CpG dinucleotide (mCG) as measured by bisulfite sequencing [35]. Earlier methods using this one-step approach include hidden Markov models (ChromHMM) [36, 37], dynamic Bayesian networks (segway) [38, 39], neural networks (CSI-ANN) [40], random forest (RFECS and REPTILE) [35, 41–43], support vector machines (SVM) [43, 44] and AdaBoost (DELTA) [45]. Although conceptually attractive and a great deal of insights into CRMs have been gained, these one-step methods have several critical limitations [35, 43, 46–51]. First, the boundaries of predicted enhancers are not well defined because of the broad enrichment of most epigenetic histone modifications in regions around CRMs, although it has been shown that using mCG as additional feature can somewhat relieve the problem [35]. Thus, the resolution of predicted CRMs is low. Second, almost all earlier methods do not predict constituent TFBSs in CRMs, in particular novel TFBSs, although it is TFBSs that largely determine the functions of CRMs. Third, many predicted CRMs cannot be validated experimentally, resulting in high FDRs [50, 51], due probably to the facts that a genome segment that has CA [15] and histone marks such as H3K4me1 [52–54], H3K4me3 [55] and H3K27ac [56] are not necessarily CRMs [46–49, 57]. Fourth, in some supervised machine-learning methods, a mark such as H3K27ac was used both as a feature and as the label of training sets [58, 59], these models therefore were actually trained to differentiate sequences with and without the H3K27ac mark, instead of the functional states of CRMs, while an active CRM may not necessarily have the mark [49, 56].

To overcome the limitations of these existing methods, we proposed a two-step approach [48, 60] (Figure 1). Specifically, in the first step, we aim to solve the CRM finding problem, i.e., given a genome, find the loci of all encoded CRMs, which is reminiscent of the earlier gene-finding problem [61], i.e., given a genome, find the loci of all encoded genes. We proposed to use all available TF ChIP-seq datasets in the organism for CRM-finding, since it has been shown that multiple TF bindings are a more reliable predictor of the loci of CRMs than CA and histone marks [46]. This is reminiscent of using all available transcripts data in the organism for gene-finding. We then predict functional states of all the putative CRMs in any cell/tissue type using few epigenetic marks in the very cell type. This goal is likely achievable, since it was found that when the locus of a CRM was accurately anchored by the bindings of multiple key TFs, epigenetic marks could be an accurate predictor of the functional state of the CRM [43, 46, 47, 51, 62]. It appears that a pattern of epigenetic marks on a CRM is sufficient to define its functional state, although a genome segment with such a pattern is not necessarily a CRM [46-48, 56, 57].

**Figure 1.**
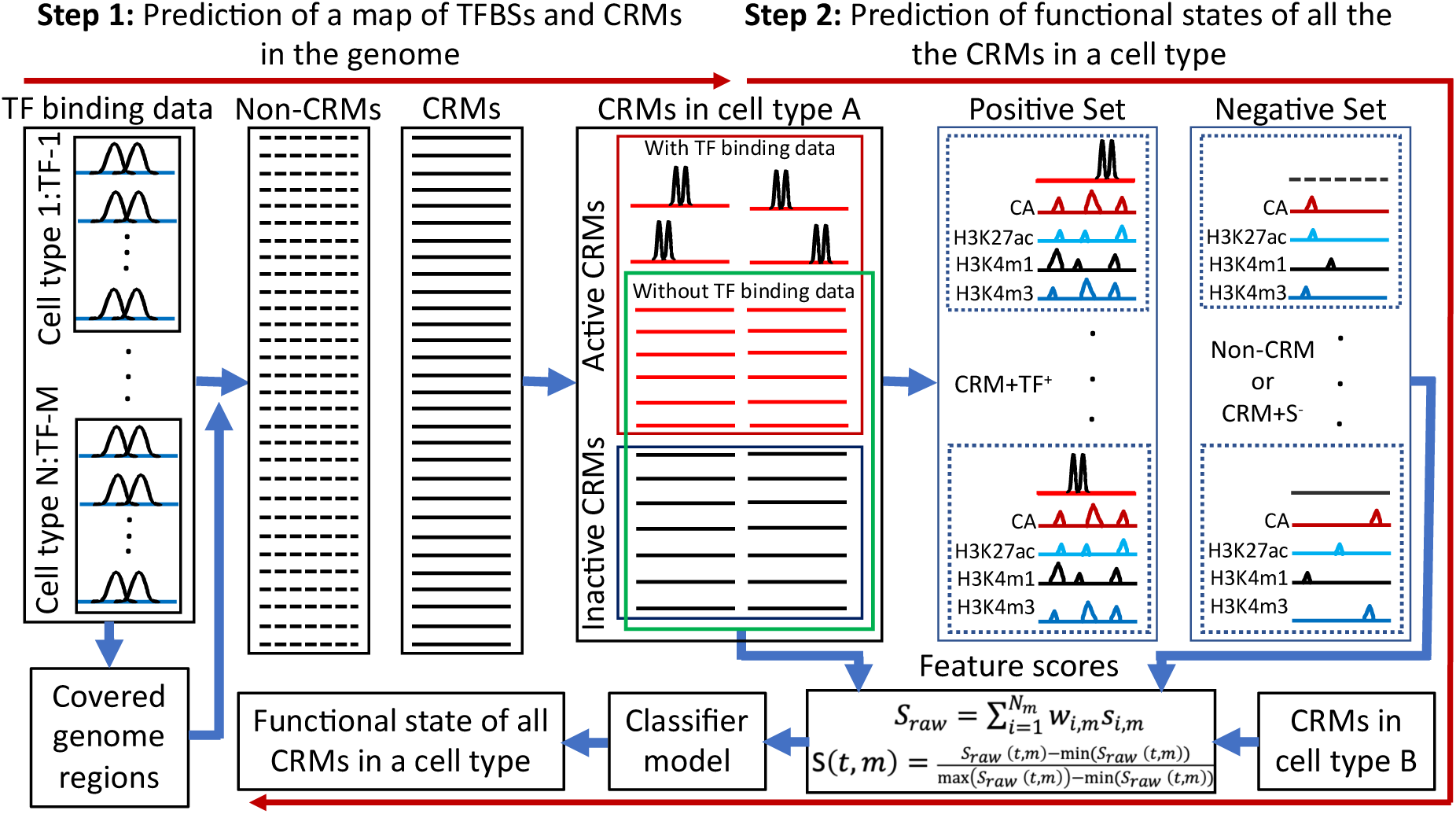
Schematic of our two-step approach and workflow of our machine-learning classifiers models. In the first step, dePCRM2 partitions the genome regions covered by all available TF binding peaks (extended to 1k-bp if shorter than 1k-bp) into a CRMs set (solid lines) and a non-CRMs set (dotted lines). In a given cell/tissue type a certain subset of the CRMs in the genome are active (red lines), while the remaining subset are non-active (black lines). We assume that a subset of the active CRMs in the cell type can be predicted by dePCRM2 based on their overlaps with available TF binding peaks in the very cell type (red lines with two binding peaks of pair-end TF ChIP-seq reads coverage). In the second step, in a cell type A with enough TF binding data, we construct a positive set (CRM+TF^+^) using the active CRMs predicted by dePCRM2 in the cell type (In most cell types, most active CRMs cannot be predicted by dePCRM2 due to insufficient availability of TF binding data in them), and a negative set either by randomly selecting predicted non-CRMs in the genome (Non-CRM), or using the putative CRMs in the genome that do not overlap STARR-seq peaks in the cell type and cannot be predicted to be active by dePCRM2 (CRM+S^-^). We train a classifier model using a few epigenetic marks on the positive and negative sets in the cell type A, or on pooled positive and negative sets from multiple cell types. The trained model is then used to predict functional states of all the CRMs whose functional states cannot be predicted by dePCRM2 in the cell type A or in any other cell type B.

For the first step, we have recently developed a pipeline dePCRM2 [48], and demonstrated its high accuracy for predicting CRMs and constituent TFBSs in the human genome using then available 6,092 TF ChIP-seq datasets for different TFs in various cell/tissue types. Nonetheless, although dePCRM2 can also predict functional states (TF-binding or non-TF-binding) in a cell/tissue type of the CRMs whose constituent TFBSs overlap binding peaks of ChIP-ed TFs in the cell/tissue type [48], the CRMs whose functional states in a cell/tissue type can be so predicted depends on the availability of TF ChIP-seq data in the very cell/tissue type. Since in most cell/tissue types, only few or even no TF ChIP-seq datasets are currently available, the fraction of CRMs whose functional state can be predicted by dePCRM2 is generally very low or even zero [48]. Obviously, to predict functional states in a cell/tissue type of all the putative CRMs in this way, one might need ChIP-seq data for all annotated TFs in the cell/tissue type. This can be too costly or currently unfeasible. Therefore, functional states of most putative CRMs in most cell/tissue types of the organism are largely unknown, and thus needed to be predicted.

In this study, we aimed to fulfill the second step of our two-step approach for predicting functional states (TF-binding or non-TF-binding) in any cell/tissue type of all the putative CRMs, particularly, those whose functional states cannot be predicted by dePCRM2 due to insufficient availability of TF binding data in the cell/tissue type (Figure 1). We showed that, using only 1∼4 epigenetic marks, machine-learning models trained on a set of CRMs whose functional states in a cell/tissue type could be predicted by dePCRM2 was able to very accurately predict functional states of the CRMs whose functional states in the cell/tissue type could not be predicted by dePCRM2. Moreover, our models using fewer epigenetic marks substantially outperform existing methods using more epigenetic marks. Thus, our two-step approach is highly accurate and cost-effective for predicting CRMs in a genome and their functional states in various cell/tissue types of the organism. Intriguingly, our models trained on cell/tissue types in human or mouse using only four epigenetic marks as the features were able to very accurately predict functional states of CRMs in developmentally distal cell/tissue types in the same and the other species. These results strongly suggest that epigenetic rules that define functional states of CRMs are common for different cell/tissue types of at least mammalian species.

## Results

### Genome-wide *de novel* prediction of CRMs in the human and mouse genomes

In our proposed two-step approach (Figure 1), we first predict a map of CRMs and constituent TFBSs in the genome using all available TF binding data in the organism. In order to predict a more complete map of CRMs in the human and mouse genomes than we did earlier using then (6/1/2019) available 6,092 and 4,786 TF binding datasets in the organisms [48, 63], we collected additional 5,256 and 4,274 TF binding-peak datasets in human and mouse cell/tissue types, respectively (Methods). The extended peaks (1,000bp) of the 11,348 (6,092+5,256, Table S1) and 9,060 (4,785+4,274, Table S2) datasets cover 85.5% and 79.9% of the human and mouse genomes, respectively. dePCRM2 [48] predicts a closely located cluster of putative TFBSs that are significantly co-occurring of multiple TF binding datasets as a CRM candidate, and a sequence between two adjacent CRM candidates in a covered genome region as a non-CRM, thereby partitioning the covered genome into two exclusive sets, i.e., the CRM candidate set and the non-CRM set, as illustrated in Figure 1. Applying dePCRM2 to these binding-peak datasets, we predicted 1,426,947 and 912,197 CRM candidates as well as 1,755,876 and 1,270,937 non-CRMs in the covered regions in the human and mouse genomes, respectively. These putative CRM candidates occupy higher proportions of the human (47.1%) and mouse (55.5%) genomes than those (44.0% and 50.4%, respectively) that we predicted earlier using smaller numbers of TF binding datasets (6,092 and 4,786, respectively). Therefore, we predicted more complete maps of CRMs and TFBSs in the genomes. Similar to our earlier predicted CRM candidates in the human genome using then available 6,092 TF ChIP-seq datasets [48], the vast majority (96%) of the predicted CRM candidate positions are located in non-coding regions in both the human and mouse genomes, and they are under either strongly positive selection (with negative phyloP scores) or strongly negative selection (with positive phyloP scores), while the predicted non-CRM positions in non-coding regions are largely selectively neutral (with near zero phyloP scores) (Figures S2A, S2B), strongly suggesting that the CRMs are likely functional, while the non-CRMs are likely not [48].

Similar to our earlier results [48], of the genome positions covered by the originally called TF binding peaks in the human or mouse genomes, only 58.7% or 75.5% were predicted to be CRM candidate positions, while the other 41.3% or 24.5% were predicted to be non-CRMs. Therefore, a called binding peak cannot be equated to a CRM. On the other hand, like our earlier results [48], of the genome positions covered by the extended parts of the originally called binding peaks, 48.7% or 58.7% were predicted to be CRM candidates, while the other 51.2% or 41.3% were predicted to be non-CRMs. The extended parts of the originally called binding peaks contribute 29.42% and 30.14% of the total predicted CRM candidate positions in the human and mouse genomes, respectively. Therefore, appropriate extension of the called short peaks can largely increase the power of the available datasets [48].

dePCRM2 [48] further evaluates each CRM candidate by computing a score and associated p-value. At a p-value cutoff of 0.05, dePCRM2 predicted 1,225,115 and 798,257 CRMs in the human and mouse genomes. We will use these putative CRMs predicted in the genomes in the remaining analysis in this study. Moreover, dePCRM2 is able to predict some putative CRMs to be active in a cell/tissue type based on overlaps between the constituent TFBSs in the CRMs and TF binding peaks available in the cell/tissue type [48]. As expected, the number of active CRMs predicted by dePCRM2 in a cell/tissue type vary widely, depending on the number of TF ChIP-seq datasets available in the cell/tissue type (Figures S1A, S1B).

### Machine-learning models trained on CRM+TF^+^/CRM+S^-^ and CRM+TF^+^/non-CRM sets outperform those trained on positive and negative sets defined by other methods

After predicting a map of CRMs in the genome of an organism, in the second step of our two-step approach (Figure 1), we predict functional states of all the putative CRMs in any cell/tissue types of the organism using few epigenetic marks by training supervised machine-learning models. Due to the lack of large gold standard sets of active CRMs and non-active CRMs in any human and mouse cell/tissue types, different methods have been used to define operational positive and negative sets of CRMs in a cell/tissue type for training machine-learning models [35, 40, 41, 45, 58, 59]. Particularly, it was recently reported [58] that STARR-seq peaks overlapping H3K27ac peaks in a cell type can be used as a high-confident set of enhancers that are active in the cell type. To find the best ways for constructing the positive and negative sets of CRMs, we evaluated seven methods (Methods, 1) in the six human cell lines (A549, HCT116, HepG2, K562, MCH-7 and SH-HY5Y) where WHG-STARR-seq data were available. Although the number of sequences in a pair of positive and negative sets is well balanced (Methods), it varies greatly in different cell/tissue types for the same method as well as for different methods due to multiple factors involved in defining the positive sets (Table 1). Particularly, as shown in Figure S1, the size of a CRM+TF^+^ set (i.e., active CRMs predicted by dePCRM2 in a cell/tissue type) depends on the number of available ChIP-seq datasets in the cell/tissue type. Moreover, since the number of predicted CRMs is much smaller than the number of 700bp-bins in the genomes, the sizes of positive sets defined based on the CRMs are generally smaller than those defined based on the genomic sequence bins (Table 1).

**Table1.** Methods for defining the seven pairs of positive and negative training sets evaluated in this study.

| Methods | Labels | Size (sequences) | Positive set | Negative set |
| --- | --- | --- | --- | --- |
| CRM+TF<sup>+</sup>/Non-CRM | TF binding | 17,558~272,128 | CRMs overlapping TF-binding peaks | Randomly selected non-CRMs |
| CRM+TF<sup>+</sup>/CRM+S<sup>-</sup> | TF binding | 17,558~272,128 | CRMs overlapping TF-binding peaks | CRMs that cannot be predicted to be active and do not overlap STARR peaks |
| CRM+S<sup>+</sup>/Non-CRM | STARR peaks | 22,610~71,176 | CRMs overlapping STARR peaks | Randomly selected non-CRMs |
| CRM+S<sup>+</sup>/CRM+S<sup>-</sup> | STARR peaks | 22,610~71,176 | CRMs overlapping STARR peaks | CRMs that cannot be predicted to be active and do not overlap STARR peaks |
| Bin+S<sup>+</sup>/Bin+S<sup>-</sup> | STARR peaks | 60,668~109,118 | 700bp bin overlapping STARR peaks | 700bp bin not overlapping STARR peaks |
| Bin+ac<sup>+</sup>/Bin+ac<sup>-</sup> | H3K27ac peaks | 175,530~1,688,868 | 700bp bin overlapping H3K27ac | 700bp bin not overlapping H3K27ac |
| Bin+S<sup>+</sup>&ac<sup>+</sup>/Bin+S<sup>-</sup>&ac<sup>-</sup> | STARR & H3K27ac peaks | 7,220~49,462 | 700bp bin overlapping STARR&H3K27ac peaks | 700bp bin not overlapping STARR&H3K27ac peaks |

We first trained a logistic regression (LR) model on each pair of the seven positive and the negative sets defined by each method (Table 1) in each of the six cell lines (A549, HCT116, HepG2, K562, MCH-7 and SH-HY5Y) using four widely available epigenetic marks (CA, H3K4me1, H3K27ac and H3K4me3) as the features (Figure 1), and validated the performance of each pair of training sets by 10-fold cross-validation. When the LR models were trained on the CRM+TF^+^/Non-CRM or CRM+TF^+^/CRM+S^-^ sets, they performed almost equally very well with a median AUROC of 0.975 and 0.976, respectively (Figure 2A).

**Figure 2.**
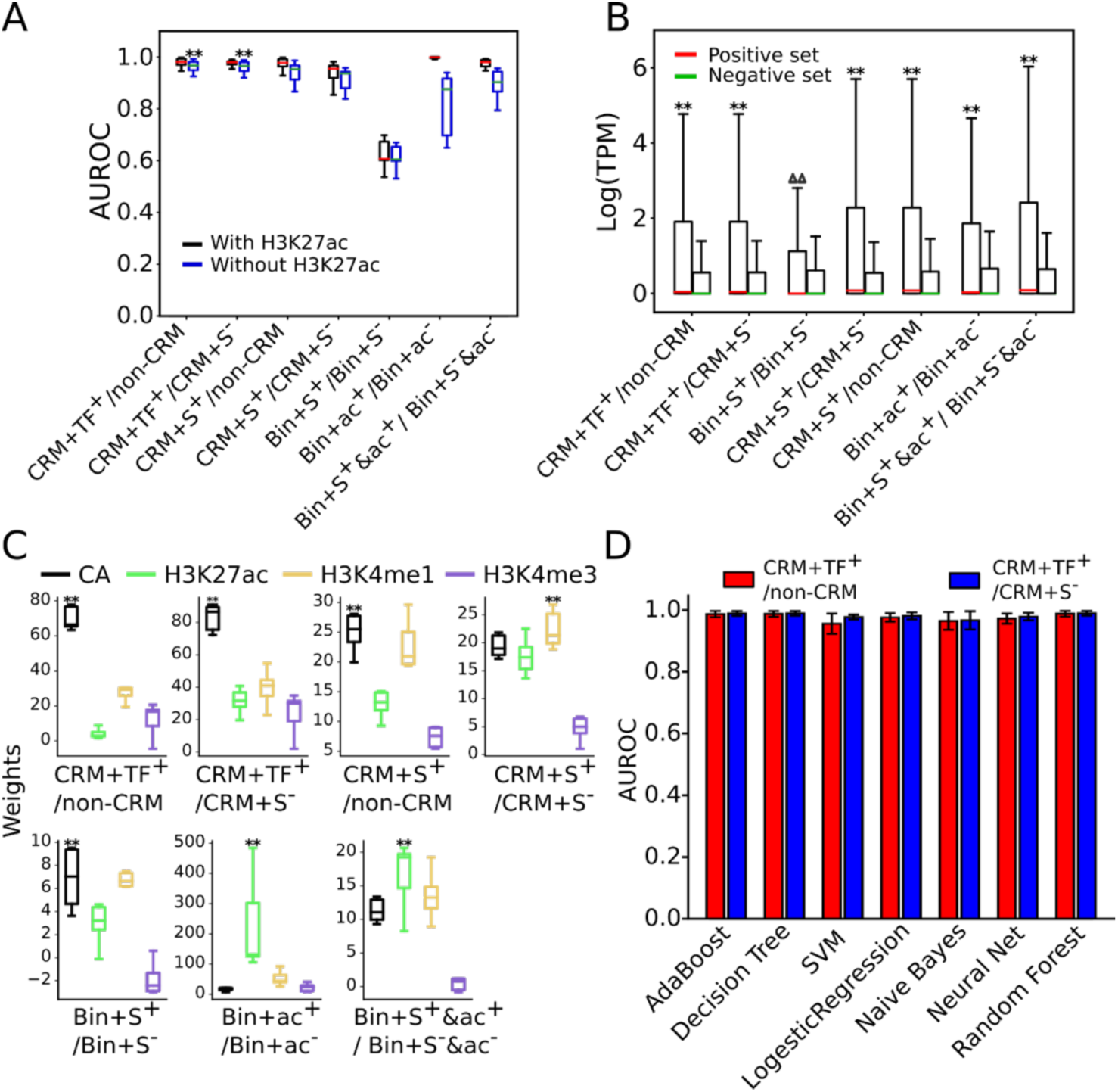
Evaluation of the seven methods for defining active and non-active CRMs. **A.** Performance of LR models trained on the pairs of positive and negative sets defined by the seven methods (Table 1) in the six cell lines using the four (CA, H3K4me1, H3K4me3 and H3K27ac) or three epigenetic marks (omitting H3K27ac) as the features. Models trained on the CRM+TF^+^/Non-CRM sets or on the CRM+TF^+^/CRM+S^-^ sets outperformed significantly those trained the other pairs of datasets when three epigenetic marks (omitting H3K27ac) were used as the features; **p<0.001 (Wilcoxon signed-rank test). **B.** Boxplot of expression levels of genes closest to the positive and negative sets defined by each method in the six cell lines. ^ΔΔ^p<4.64x10^-56^, **p<2.2x10^-302^ (Mann Whitney U test). **C.** Boxplot of weights (coefficients) of the four epigenetic marks in the LR models for discriminating the positive and negative sets defined by each method in the six cell lines. **p<0.001 (Wilcoxon signed-rank test), the weights of the marks are significantly higher than those on the other three marks. **D.** Performance of the seven types of classifiers trained on the CRM+TF^+^/Non-CRM sets or on the CRM+TF^+^/CRM+S^-^ sets in the six cell lines using the four epigenetic marks as the features.

Consistently, the positive set CRM+TF^+^ had distinct patterns of the four marks from both the negative sets, i.e., Non-CRM (Figure S3A) and CRM+S^-^ (Figure S3B). Interestingly, the two negative sets had similarly low levels of the four marks (Figure S3C), suggesting that the CRM+S^-^ set are indeed non-active and that non-active CRMs in a cell/tissue type might also have highly similarly low levels of the four marks. Distributions of phyloP scores [64] indicates that non-coding positions of both the CRM+TF^+^ sets (Figure S4A) and the CRM+S^-^ sets (Figure S4B) are subject to substantially more evolutionarily constrained than those of the entire 85.5% genome regions covered by the extended TF binding peaks, and thus might be true CRM loci as we argued earlier [48]. Nonetheless, it is not surprising that the CRM+TF^+^ sets and the CRM+S^-^ sets differ in their functional states, as they have very differed in epigenetic mark patterns (Figure S3B). In stark contrast, non-coding positions of the negative sets Non-CRM are more likely selectively neutral than those in the entire covered genome regions (Figure S4A), and are unlikely CRMs as we argued earlier [48].

Consistently, genes closest to the CRM+TF^+^ sets had significantly higher expression levels (p<2.23x10^-302^) than those closest to the CRM+S^-^ or Non-CRM sets, and the latter two groups of genes had very low expression levels (Figure 2B). Taken together, these results suggest that the CRM+TF^+^ sequences are indeed active CRMs in the cell line and the CRM+S^-^ sequences are CRMs but are not active in the cell line, while Non-CRM sequences are even not CRMs.

When trained on the CRM+S^+^/non-CRM sets and the CRM+S^+^/CRM+S^-^ sets, the models also performed well with a median AUROC of 0.972 and 0.943 (Figure 2A), respectively, albeit slightly worse than those trained on the CRM+TF^+^/Non-CRM sets (median AUROC=0.975) or on the CRM+TF^+^/CRM+S^-^ sets (median AUROC=0.976) (Figure 2A). Consistently, the positive sets CRM+S^+^ have distinct patterns of the four epigenetic marks from those on the two negative sets Non-CRM (Figure S3D) and CRM+S^-^ (Figure S3E). As expected, non-coding positions of the CRM+S^+^/Non-CRM sets (Figure S4C) and of the CRM+S^+^/CRM+S^-^ sets (Figure S4D) evolve quite similarly to those of the CRM+TF^+^/Non-CRM sets (Figure S4A) and those of the CRM+TF^+^/CRM+S^-^ sets (Figure S4B), respectively. In agreement, genes closest to the positive sets CRM+S^+^ had significantly higher expression levels (p<2.23x10^-302^) than those closest to the negative sets CRM+S^-^ or Non-CRM sets (Figure 2B). Taken together, these results suggest that at least most of the CRM+S^+^ sequences are indeed active CRMs in the cell line.

To our surprise, the models trained on the Bin+S^+^/Bin+S^-^ sets only achieved a mediocre median AUROC of 0.621 (Figure 2A), indicating that the models had little capability to discriminate genome sequence bins that overlapped STARR-seq peaks (Bin+S^+^) and those that did not (Bin+S^-^). Consistently, the two sets had little differences in their patterns of the four epigenetic marks (Figure S3F). Moreover, as shown in Figure S4E, non-coding positions in both sets evolve much like non-coding positions of the entire covered regions of the human genome [48], suggesting that like the negative sets Bin+S^-^, a large portion of the positive sets Bin+S^+^ were not even CRMs. In agreement, genes closest to the Bin+S^+^ sets had very low expression levels, which were only slightly, yet significantly higher (p<4.64x10^-58^) than those closest to the Bin+S^-^ sets (Figure 2B). Such low expression levels of the genes are understandable, as it has been shown that episomal expression vectors used to define WHG-STARR-seq peaks do not mimic native chromosomal contexts of assessed sequences, resulting in up to 87.3% FDR [17].

When trained on the Bin+ac^+^/Bin+ac^-^ sets or on the Bin+S^+^&ac^+^/Bin+S^-^&ac^-^ sets, the models could very accurately discriminate the positive sets and the negative sets with a very high median AUROC of 0.996 and 0.973, respectively (Figure 2A), in agreement with the earlier reports [58, 59] that used similarly constructed positive and negative sets. Consistently, the positive sets and negative sets defined by both methods had distinguishable patterns of the four epigenetic marks (Figures S3G, S3H). However, we suspected that the superior performance on both pairs of training sets might be artifacts due to the aforementioned reason that H3K27ac was used both as one of the features and as the label of the training sets in both methods (Table 1). Indeed, when the models were trained on the Bin+ac^+^/Bin+ac^-^ sets or the Bin+S^+^&ac^+^/Bin+S^-^&ac^-^ sets using only the other three marks (CA, H3K4me1 and H3K4me3) as the features, their performance reduced by 17.1% and 8.0% with an intermediate median AUROC of 0.826 and 0.895, respectively (Figure 2A). Although non-coding positions of both positive sets Bin+ac^+^ and Bin+S^+^&ac^+^ are under moderately more evolutionary constraints than those in the respective negative sets Bin+ac^-^ and Bin+S^-^&ac^-^, their phyloP score distributions differ only slightly from those of the entire covered regions of the genome (Figures S4F, S4G), suggesting that a considerable portion of the positive sets defined by both methods might not be even CRMs, although they were heavily marked by H3K27ac (Figures S3G, S3H) and genes closest to Bin+ac^+^ and Bin+S^+^&ac^+^ sets had higher expression levels (p<2.23x10^-302^) than tho**s**e closest to Bin+ac^-^ and Bin+S^-^&ac^-^ sets, respectively (Figure 2B). Since sequences marked by H3K27ac are not necessarily CRMs [46, 47], and H3K27ac is not essential for active CRMs [56], it appears that the models were trained to differentiate genomic sequence bins that were marked by H3K27ac or not, rather than active and non-active CRMs.

Unlike cases of the Bin+ac^+^/Bin+ac^-^ and Bin+S^+^&ac^+^/Bin+S^-^&ac^-^ sets that used H3K27ac as the label, omitting H3K27ac as one of the features had little effects on the performance of the models trained on all the other five pairs of positive and negative sets that did not use H3K27ac as the label (Figure 2A, Table 1). Notably, when H3K27ac was omitted as a feature, models trained on the CRM+TF^+^/non-CRM sets and the CRM+TF^+^/CRM+S^-^ sets performed significantly better (p<0.001) than those trained on all the other pairs of datasets (Figure 2A, Table 1). Consistently, H3K27ac had the highest weights (p<0.001) in the models trained on the Bin+ac^+^/Bin+ac^-^ and Bin+S^+^&ac^+^/Bin+S^-^&ac^-^ sets, while CA had the highest weights (p<0.001) in the models trained on the other five pairs of datasets except for the CRM+S^+^/CRM+S^-^ sets where H3K4me1 had the highest weights (p<0.001) (Figure 2B). These results indicate that a mark should not be used both as a feature and as the label in training datasets to avoid artifacts.

Notably, models trained on the positive set CRM+TF^+^ perform better than those trained on the positive set CRM+S^+^ no matter whether the Non-CRM set or the CRM+S^-^ set was used as the negative set (Figure 2A). To reveal the subtle differences in their epigenetic modifications, we plotted heat maps of signals of the four epigenetic marks around each positive set (CRM+TF^+^ or CRM+S^+^) and its two size-matched negative sets (non-CRM and CRM+S^-^) in a cell line. As shown in Figures 3, S5, in all the cell lines (raw data in SH-HY5Y cells were not available to us), the two negative sets indeed had virtually indistinguishable patterns of the four marks as indicated earlier (Figure S3C), while the positive set CRM+TF^+^ had stronger CA and H3K4me3 signals than the positive set CRM+S^+^, and the reverse was true for the H3K27ac and H3K4me1 signals. The stronger CA signals of the CRM+TF^+^ set might largely account for the better performance of models trained on it than those trained on the CRM+S^+^ set, given that CA was the most important feature in the models (Figure 2C). Taken together, these results suggest that the putative CRMs that are predicted by dePCRM2 to be active in a cell/tissue type (i.e., CRM+TF^+^) are more likely to be active than our putative CRMs that overlap STARR-seq peaks in the cell type (CRM+S^+^). Consistent with this conclusion, it was reported that not all STARR-seq peaks would be active in the native chromatin environment, as the method quantifies enhancer activity in an episomal fashion [15–17, 19]. In summary, the LR models trained on the CRM+TF^+^/Non-CRM sets and CRM+TF^+^/CRM+S^-^ sets perform equally well, and they substantially outperform models trained on the other five pairs of positive/negative sets.

**Figure 3.**
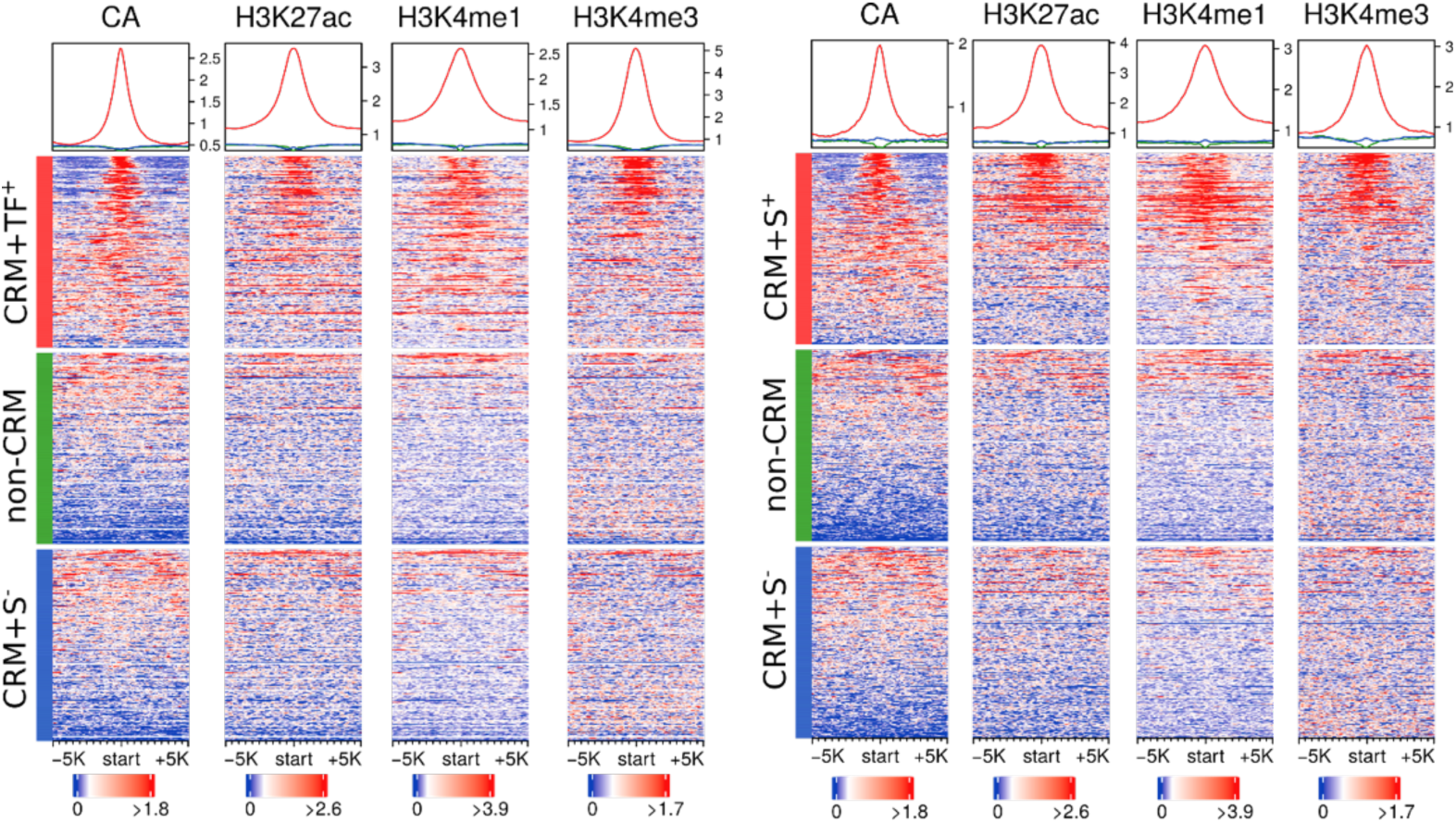
Heat maps of signals of the four epigenetic marks around the positive sets CRM+TF^+^ and CRM+S^+^, and the negative sets non-CRM and CRM+S^-^ in the HCT116 cells. We plotted signals of CA, H3K27ac, H3K4me1, and H3K4me3 around the sequences in each set. To make the density plots, we first extended the centers of sequences to 10 kb, and then for each 100-bp tiling window in the extended regions, we calculated the signal scores using EnrichedHeatmap (w0 mode) [65]. Note that the scales of vertical axes of the density plots on the top of heat maps are different in the right and left panels. The sequences in each set are sorted in the descending order of the CA signals for all the marks.

We next asked how six other machine-learning classifiers (AdaBoost, SVM, neural network, naïve Bayes, decision tree, random forest) perform when their models are trained on the CRM+TF^+^/Non-CRM sets and CRM+TF^+^/CRM+S^-^ sets in the six cell lines using the four marks as the features. As shown in Figure 2D, like LR models, models of all these six classifiers trained either on the CRM+TF^+^/Non-CRM sets or on the CRM+TF^+^/CRM+S^-^ sets also achieved a very high median AUROC (>0.970), indicating that both pairs of positive and negative sets can be accurately and robustly differentiated, presumably due to the distinct patterns of the four epigenetic marks on the two positive sets and the two negative sets (Figures 3, S3A, S3B, S5). Notably, random forest, decision tree and AdaBoost slightly outperformed the other four classifiers including LR. However, we chose LR for further analysis, as the weights (coefficients) in the models are consistent with those of the linear SVM models (data not shown), and are more explainable. Moreover, models of all the classifiers trained on the CRM+TF^+^/CRM+S^-^ sets slightly outperformed those trained on the CRM+TF^+^/Non-CRM sets (Figure 2D). Using the CRM+S^-^ sequences as the negative set also logically makes more sense than using the Non-CRM sequences. Nonetheless, since STARR-seq data were available in only few cell lines, and since the Non-CRM and CRM+S^-^ sets in a cell types had virtually indistinguishable patterns of the four epigenetic marks (Figures 3, S3C and S5), we used the CRM+TF^+^/Non-CRM sets as the training sets in the remaining predictions and analyses in the 67 human and 64 mouse cell/tissue types that have datasets available of the four epigenetic marks (Methods).

### Few epigenetic marks are sufficient to accurately predict functional states of the CRMs

It was recently reported that machine-learning models trained using six epigenetic marks (H3K27ac, H3K4me1, H3K4me2, H3K4me3, H3K9ac and CA) had almost the same power as models trained using 30 marks in differentiating STARR-seq peaks overlapping H3K27ac peaks and negative control sequences [58]. We thus asked whether an even smaller set of epigenetic marks in a cell/tissue type can accurately predict functional states of all the putative CRMs in the genome of the organism, respectively, particularly of those CRMs whose functional states cannot be predicted by dePCRM2 due to the lack of sufficient TF-binding data in the cell/tissue type. To this end, we first evaluated the 15 possible combinations of the four arguably most important epigenetic marks (CA, H3K4me1, H3K27ac and H3K4me3) that have widely available data. Using each of the 15 combinations of the four marks as the features, we trained a LR model on the CRM+TF^+^/Non-CRM sets in each of the 67 human and 64 mouse cell/tissue types, and evaluated the model using 10-fold cross-validation. As shown in Figure 4A, in a human cell/tissue type, H3K27ac peaks on average covered the largest number of genome positions, followed by H3K4me1, H3K4me3 and CA peaks. Although there were on average extensive overlaps between each pair of the four marks in a cell/tissue type, only a small number of genome positions were covered by peaks of all the four marks. For each mark, its unique genome positions were the most predominant form among all possible ways that it overlapped other marks (Figure 4A).

**Figure 4.**
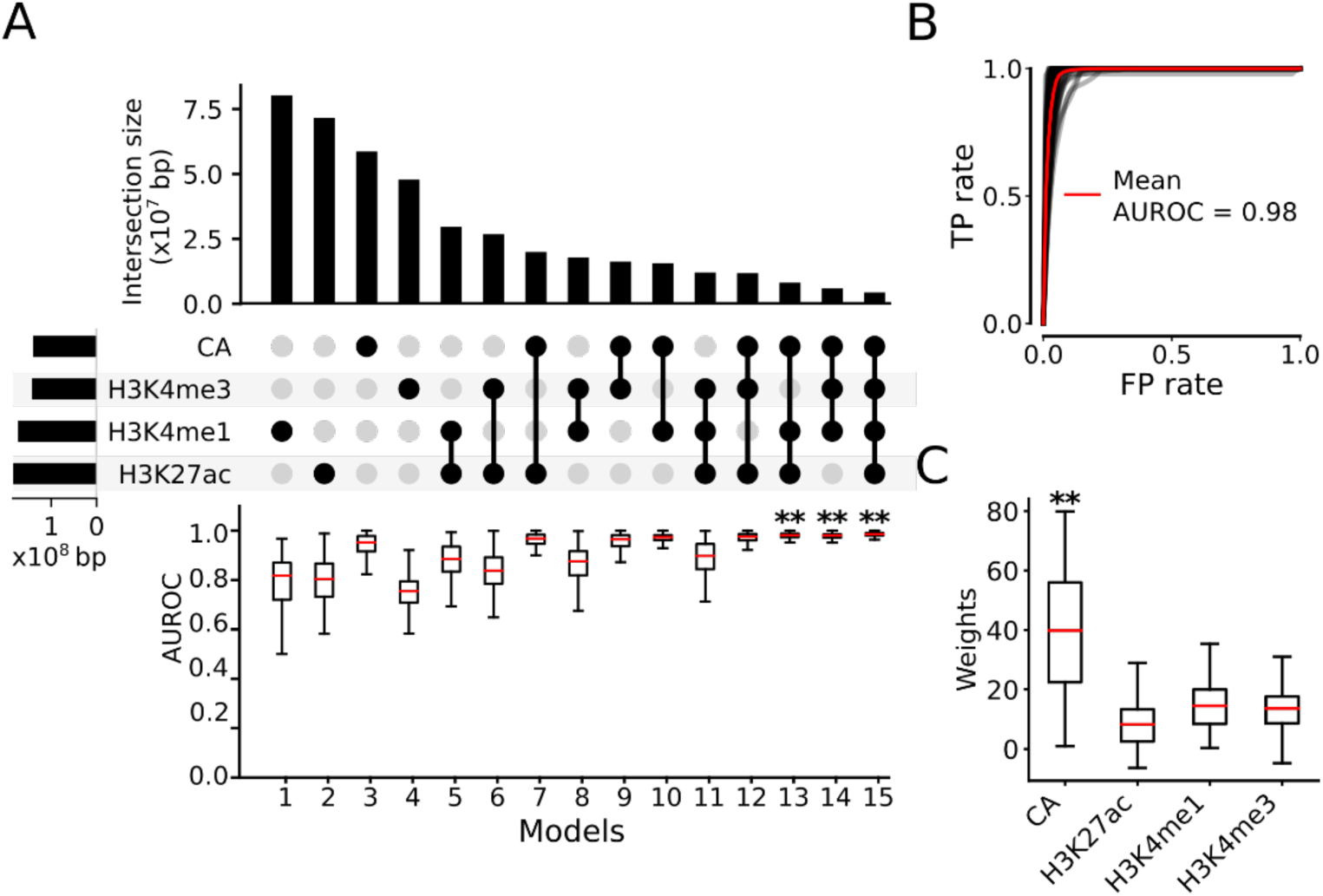
Identification of optimal minimal sets of epigenetic marks for predicting functional states of CRMs in human cell/tissue types. **A.** Upset plot showing the mean unique genome positions of the peaks of each mark and the mean intersections of genome positions covered by the peaks of the four marks (upper and middle panels), and boxplot of AUROCs of the 15 LR models using the 15 different combinations of the four marks in the 67 human cell/tissue types. Models 13, 14 and 15 have similar AUROCs, each is significantly higher than those of the other 12 models, **p<0.001 (Wilcoxon signed-rank test) **B.** ROC curves of model 15 using the four marks in the human cell/tissue types. Each gray curve is the ROC curve for a cell/tissue type, and the red one is the mean ROC curve of the 67 cell/tissue types. **C.** Boxplot of weights (coefficients) of the four marks in model 15. **p<0.001 (Wilcoxon signed-rank test), the weights of CA are significantly higher than those of the other three marks.

Each single mark on average had varying capability of differentiating the CRM+TF^+^/Non-CRM sets, with CA (model 3) performing best, followed by H3K4me1 (model 1), H3K27ac (model 2), and H3K4me3 (model 4), with a median AUROC of 0.953, 0.818, 0.803, and 0.755, respectively (Figure 4A). This result is in excellent agreement with the earlier finding that CA alone can be a good predictor of activities of CRMs when combined with multiple TF binding data [51]. However, we noted that the models for each single mark could perform poorly in some cell/tissue types (Figure S6), due probably to the low quality of the datasets collected from them. Among the six models using a combination of two marks as the features, model 10 using CA+H3K4me1 achieved the highest median AUROC of 0.974 (Figure 4A). Of the four models using a combination of three marks as the features, model 13 using CA+H3K4me1+H3K27ac (median AUROC=0.981) and model 14 using CA+H3K4me1+H3K4me3 (median AUROC=0.980) outperformed the other two models (p<0.001) (Figure 4A). Model 15 using all the four marks (CA+H3K4me1+H3K27ac+ H3K4me3) had the highest median AUROC of 0.986 among all the 15 models (p<0.001) (Figures 4A and 4B). CA had the highest contribution among the four marks to predicting functional states of CRMs in model 15 (Figure 4C), in agreement with the earlier result based on the six cell lines (Figure 2C). This is in stark contrast with the earlier result that H3K27ac was the most important feature for predicting H3K27ac labelled sequences [58], due probably to the aforementioned reason that the mark was used both as a feature and as the label (Figures 2A and 2C). Interestingly, with the increase in the number of marks used as the features, the variation of performance of the models in different cell/tissue types decreased (Figure S6), suggesting that the effects of a low-quality dataset for a mark can be compensated by using datasets of other marks in the cell/tissue type.

As shown in Figure 4A, adding an additional mark to the feature list of a model always led to a new one that outperformed the original one. However, the infinitesimal increments of 0.005 (0.5%) and 0.006 (0.6%) in the median AUROC of model 15 over models 13 (CA+H3K4me1+H3K27ac) (0.981) and 14 (CA+H3K4me1+ H3K4me3) (0.980) (Figure 4A), respectively, suggest that the improvement of accuracy is already in the later phase of saturation. To verify this, we trained LR models using five (adding H3K4me2 or H3K9ac to the four marks) or six (adding both H3K4me2 and H3K9ac to the four marks) marks as the features on 22 human cell/tissue types in which all the six marks datasets were available (Table S3). As shown in Figure S7A, the models trained on the four marks achieved a median AUROC of 0.9696 in the 22 human cell/tissue types, while adding H3K9ac or H3K4me2 to the four marks improved the median AUROC (0.9701 or 0.9710) by only 0.0005 (0.05%) or 0.0014 (0.14%), respectively, and adding both marks improved the median AUROC (0.9718) by only 0.0022 (0.22%). Thus, improvement of accuracy by using more than four marks is indeed very limited. Therefore, four marks (CA+H3K4me1+ H3K4me3+H3K27ac), optimal combinations of three marks (CA+H3K4me1+H3K27ac, or CA+H3K4me1+H3K4me3), optimal combination of two marks (CA+H3K4me1), or even single mark (i.e., CA) is sufficient to very accurately predict functional states of our putative CRMs in a human cell/tissue type, though the more marks used as the features, the more accurate prediction obtained (Figure 4A). In the model using the six marks as the features, CA again had the highest weights, followed by H3K4me1, H3K4me2, H3K4me3, H3K27ac and H3K9ac (Figure S7B).

The rapid saturation of AUROC values with the increase in the number of marks used as the features suggests redundancy of information in data of different marks. To reveal this, we computed Pearson’s correlation coefficients (γ) between each pair of the six marks on the CRMs in the positive set (CRM+TF+) that were predicted by dePCRM2 to be active in each of the 22 human cell/tissue types. Indeed, all pairs of the six marks have varying levels of positive correlations (Figure S7C). Specifically, H3K4me1 and H3K9ac (γ=0.12) as well as H3K4me1 and H3K4me3 (γ=0.14) have low correlations; H3K4me3 and H3K9ac (γ=0.86) as well as H3K27ac and H3K9ac (γ=0.81) have high correlations; and all the other pairs have intermediate (γ=0.27∼0.73) correlations. The correlations can also be seen from heat maps of signals of the six marks around the positive sets (CRM+TF^+^) in each cell/tissue type as shown in Figure S7D for the GM12878 cells as an example.

The same conclusions are drawn from the results obtained using the mouse datasets (Figures S8∼S10). However, the models generally performed better in the mouse datasets (Figures S8∼S10) than in the human datasets (Figures 4, S6, S7), due probably to the better quality of mouse datasets, as evidenced by the less variation of the performance of the models using single marks (Figure S9 vs Figure S6). Therefore, we used the four marks (CA+H3K4me1+H3K4me3+H3K27ac) as the features in the remaining predictions and analyses, considering the wider availability of their data, albeit H3K4me2 had slightly higher weights in the models than H3K4me3, H3K27ac and H3K9ac (Figures S7B, S10B).

### Epigenetic rules defining functional states of CRMs are the same in different cell types in human or mouse and even across human and mouse

It has been shown that when mCG is used along with other six epigenetic marks (H3K4me1, H3K4me2, H3K4me3, H3K27me3, H3K9ac, H3K27ac), machine-learning models trained on mouse embryonic stem cells (mESCs) or human H1 embryonic stem cells (H1-hESCs) could accurately predict active enhancers in other developmentally closely related mouse and human cell/tissue types [35]. To see whether distinct epigenetic patterns of the four epigenetic marks (CA, H3K4me1, H3K4me3, and H3K27ac) on our positive sets CRM+TF^+^ and negative sets Non-CRM (Figures 3, S3A, S5) learned in many cell/tissue types in a species can be transferred to any other cell/tissue types in the same species, we trained LR models on pooled positive and negative sets from n-1 human cell/tissue types (n=67), and tested it on the left-out cell/tissue type (leave-one-out cross-validation, see Methods). As shown in Figure 5A, the models trained on different cell/tissue types achieved quite a high median AUROC of 0.985 in the left-out ones, which was not significantly different from that (0.986) achieved by the models that were trained and tested on the same cell/tissue types using 10-fold cross-validation (p<0.327). Similar results (median AUROC= 0.990 vs. 0.990) were obtained using the datasets from 64 mouse cell/tissue types (Figure 5B, p<0.353). Given the highly developmentally distal nature of these human (Table S3) and mouse (Table S4) cell/tissue types, these results strongly suggest that the rules that define active and non-active CRMs might be the same in even developmentally distal cell/tissue types in the same species.

**Figure 5.**
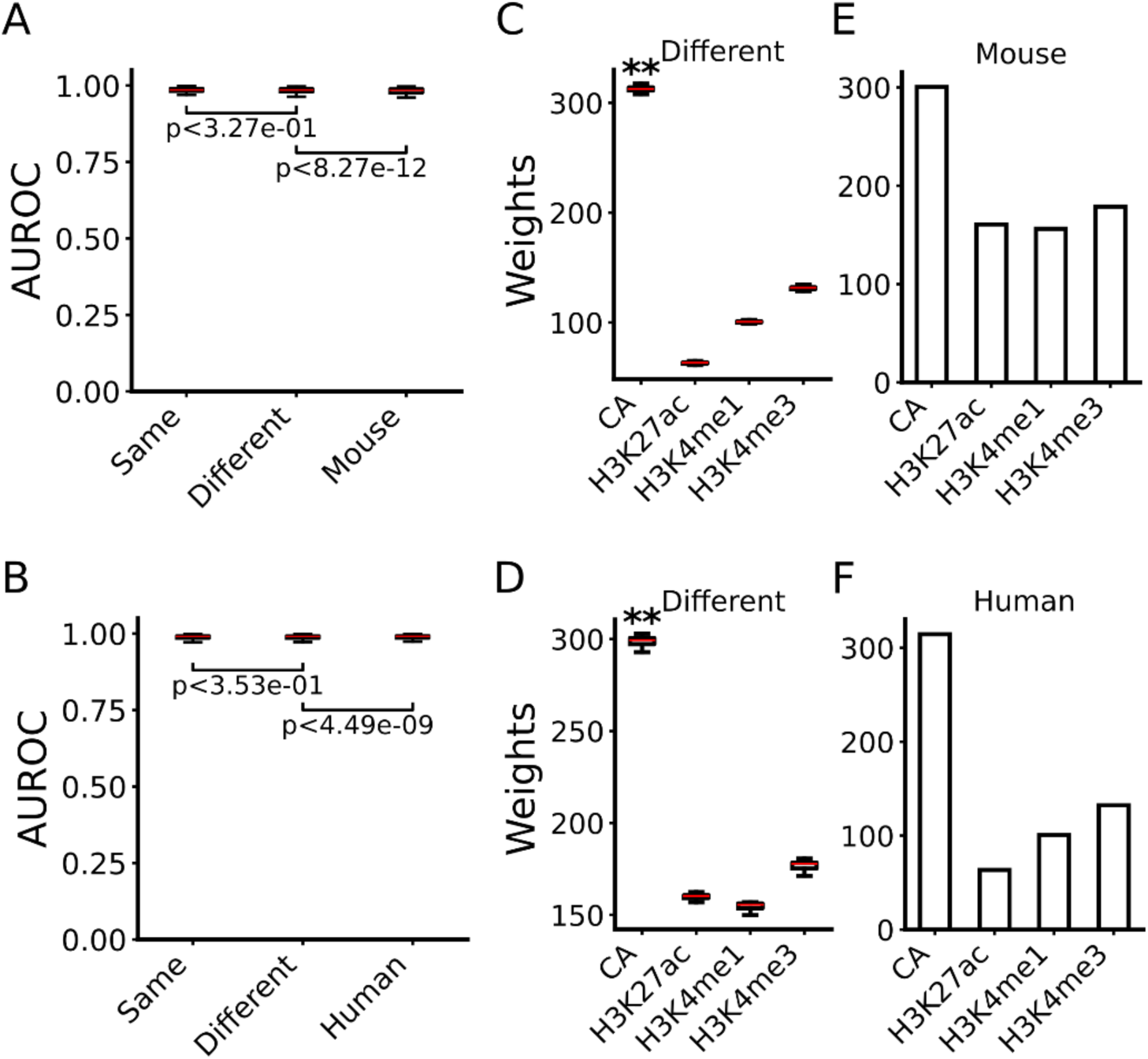
Common epigenetic rules defining functional states of CRMs in various cell/tissue types of the same and different species. **A, B.** Boxplots of AUROCs of the models trained and tested in the same cell/tissue types in human (A) or mouse (B), of the models trained in multiple cell/tissue types in human (A) or mouse (B) and tested in different cell/tissue types in the same species, and of the model trained in multiple cell/tissue types in mouse (A) or human (B) and tested in various cell/tissue types in human or in mouse, respectively. The statistical tests were done using Wilcoxon signed-rank test. **C, D.** Boxplots of coefficients of the four marks in the LR models trained in multiple cell/tissue types in human (C) or mouse (D) and tested in different cell/tissue types in the same species. **p<0.001 (Wilcoxon signed-rank test), the weights of CA are significantly higher than those of the other three marks. **E, F.** Coefficients of the four marks in the model trained in multiple cell/tissue types of mouse (E) or human (F) and tested in various cell/tissue types of human or mouse, respectively.

Moreover, in a more recent study, it was found that that machine-learning model trained on *Drosophila* S2 cells using six epigenetic marks (CA, H3K9ac, H3K27ac, H3K4m1,H3K4m2, and H3K4m3) as the features could be transferred to predict active promoters and enhancers in mouse and human cell/tissues [58]. To see whether patterns of the four epigenetic marks on active and non-active CRMs learned in multiple cell/tissue types of a mammal can be transferred to various cell/tissue types of another mammal, we trained LR models on pooled training datasets from the 64 mouse cell/tissue types, and tested it on each of the 67 human cell/tissue types, and vice versa. As shown in Figures 5A and 5B, very high median AUROCs of 0.984 and 0.991 were achieved in the human and mouse cell types using the models trained on the mouse and human cell/tissue types, respectively, which were only slightly, though significantly different (p<8.27x10^-12^ and p<4.49x10^-9^) from those achieved by the models that were trained and tested in the same species (median AUROC=0.985 and 0.990), respectively. These results strongly suggest that epigenetic patterns that define active and non-active CRMs are highly conserved in different cell/tissue types of humans and mice. As in the earlier case where the models were trained and tested in the same cell/tissue types (Figures 4C, S8C), CA was the most important feature for predicting functional states of CRMs in these two latter cases (Figures 5C∼5F). Therefore, it appears that epigenetic rules defining functional states of CRMs are the same in different cell/tissue types of different mammalian species, though only two species (humans and mice) were tested here.

### The models have similar performance in predicting proximal and distal CRMs

CRMs can be largely classified into proximal and distal ones based on their distances to their nearest transcription start sites (TSSs). The former category often overlaps TSSs and functions as core or proximal promoters, while the latter category often functions as enhancers or silencers, though such classification is not clear-cut, as some promoters can also function as distal enhancers of remote genes [66]. The distances between our predicted CRM and their nearest TSSs show a bimodal distribution in both the human (Figure S11A) and mouse (Figure S11B) genomes. The proximal CRMs (distance to the nearest TSS ≤ 1,000bp) make up 10.5% and 9.5% of the putative CRMs in the human and mouse genomes, while the remaining 89.5% and 91.5% of the CRMs are distal, respectively. To see how well the LR models perform for predicting these two categories of CRMs, we split both a positive set and a negative set into a proximal set and a distal set, and evaluated the models using these split proximal and distal sets. We found that models trained on the CRM+TF+/Non-CRM sets in a cell/tissue type achieved similarly high accuracy for predicting active proximal and distal CRMs in the same cell/tissue types in both human (0.989 and 0.983, respectively) and mouse (0.992 and 0.988, respectively) cell/tissue types (data not shown).

To further verify this and to see whether active proximal and distal CRMs have distinct epigenetic mark patterns, we separately trained LR models on the split proximal and distal positive and negative sets. When trained and tested in the same cell/tissue types using 10-fold cross-validation, the models performed well for predicting active proximal and distal CRMs in both human and mouse cell/tissue types, with a median AUROC of 0.989 and 0.983 (Figures 6A, 6B), and 0.992 and 0.988 (Figures 7A, 7B), respectively. The models trained on n-1 cell/tissue types in human and mouse also performed well in left-out cell types in the same species in predicting active proximal and active distal CRMs, with a median AUROC of 0.988 and 0.984 (Figures 6A, 6B), and 0.993 and 0.988 (Figures 7A, 7B), respectively, which were not significantly different from or even better than those of models trained and tested in the same cell/tissue types (Figures 6A,6B, 7A, 7B), due probably to the larger training stets (n-1 vs 1). The models trained on multiple mouse or human cell/tissue types also performed well in various cell/tissue types in the other species in predicting active proximal (Figures 6A, 6B) and active distal CRMs (Figures 7A, 7B), with a median AUROC of 0.988 and 0.982, and 0.993 and 0.989, respectively. Thus, the performance of the models was only slightly though significantly (p<2.23x10^-11^ and p<5.87x10^-11^) different from that of the models trained and tested in the same species. Interestingly, in all the models of the three scenarios in both human and mouse, CA and H3K4me3 contributed more (p<0.001) than H3K4me1 and H3K27ac for predicting active proximal CRMs (Figures 6C∼6H), while CA and H3K4me1 contributed more (p<0.001) than H3K4me3 and H3K27ac for predicting active distal CRMs (Figures 7C∼7H). These results are consistent with the findings that both active promoters and active enhancers are nucleosome free [23–25], but the former tend to have a higher level of H3K4me3 and a lower level of H3K4me1 in flanking regions, and the opposite is true for the latter [54, 55, 67–71].

**Figure 6.**
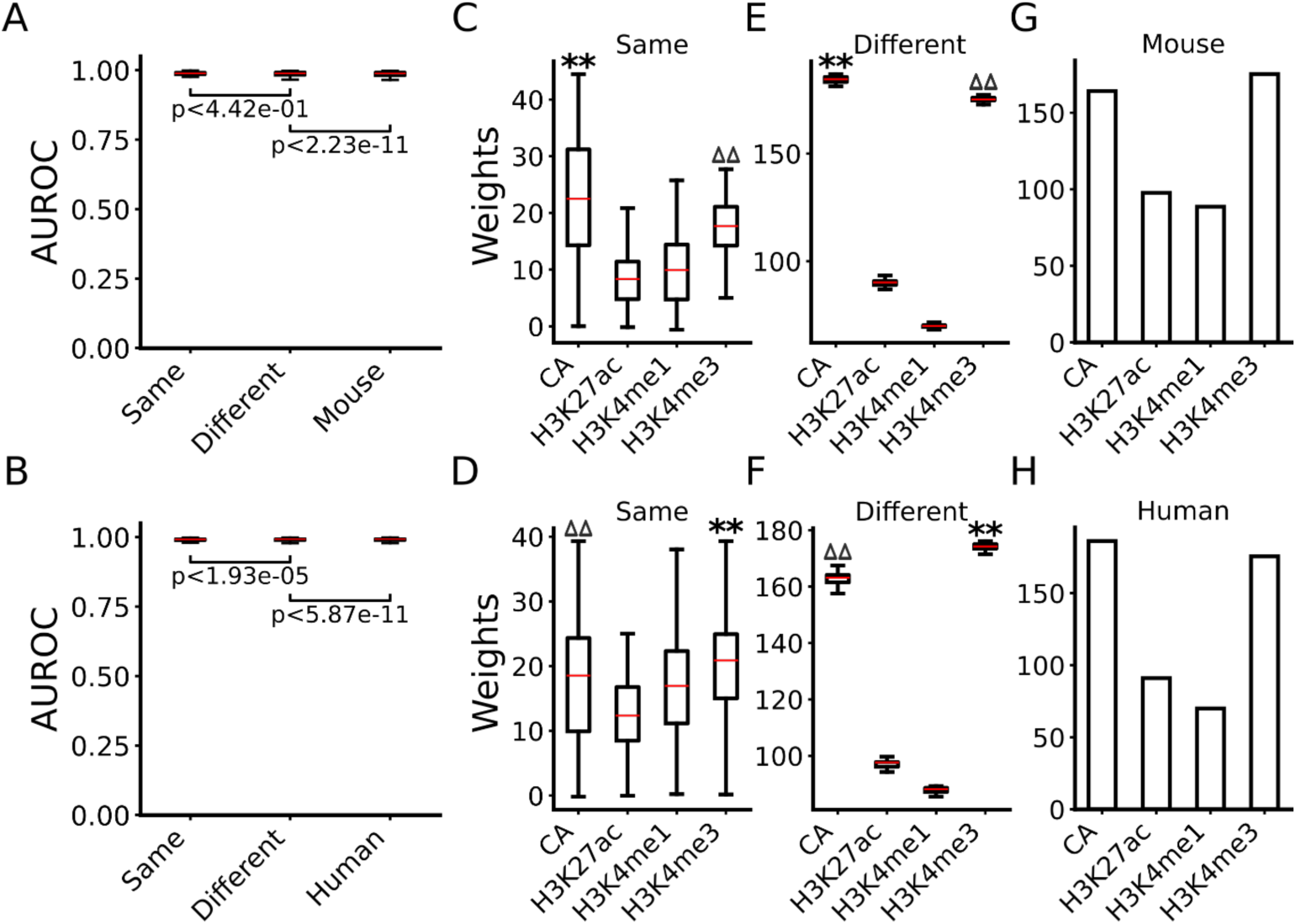
Performance of the models for predicting the functional states of proximal CRMs. **A, B.** Boxplots of AUROCs of the models trained and tested in the same cell/tissue types in human (A) or mouse (B), of the models trained in multiple cell/tissue types in human (A) or mouse (B) and tested in different cell/tissue types in the same species, and of the model trained in multiple cell/tissue types in mouse (A) or human (B) and tested in various cell/tissue types in human or mouse, respectively. The statistical tests were done using Wilcoxon signed-rank test. **C, D.** Boxplots of coefficients of the four marks in the models trained and tested in the same cell/tissue types in human (C) or mouse (D). **E, F.** Boxplots of coefficients of the four marks in the models trained in multiple cell/tissue types in human (E) or in mouse (F) and tested in different cell/tissue types in the same species. **p<0.001 (Wilcoxon signed-rank test), the weights of the labeled mark are significantly higher than those of the other three marks; ^ΔΔ^p<0.001 (Wilcoxon signed-rank test), the weights of the labeled mark are significantly higher than those of the other two marks. **G, H.** Coefficients of the four marks in the model trained in multiple cell/tissue types in mouse (G) or human (H) and tested in various cell/tissue types in human or mouse, respectively.

**Figure 7.**
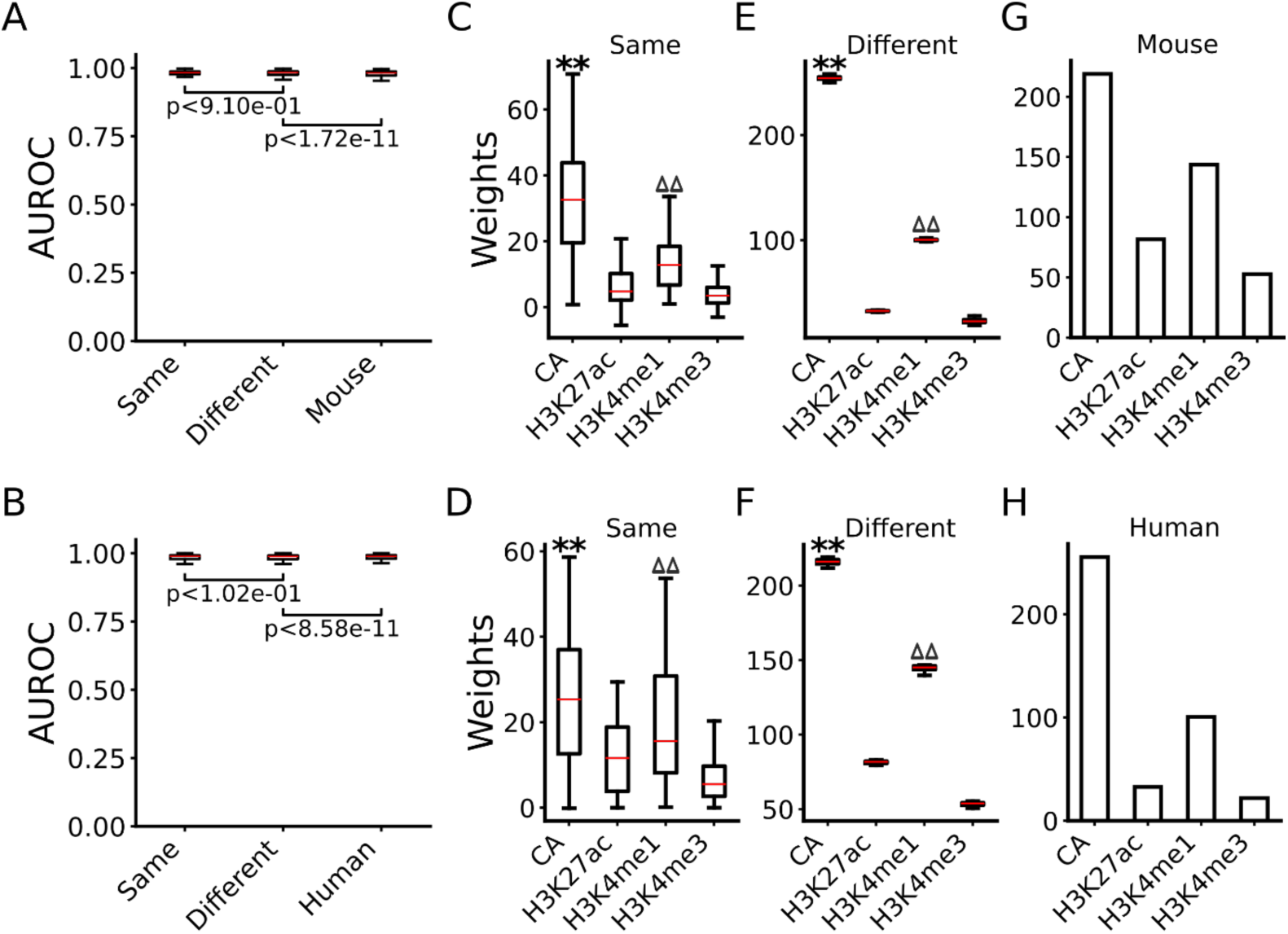
Performance of the models for predicting the functional states of distal CRMs. **A, B.** Boxplots of AUROCs of the models trained and tested in the same cell/tissue types in human (A) or mouse (B), of the models trained in multiple cell/tissue types in human (A) or mouse (B) and tested in different cell/tissue types in the same species, and of the model trained in multiple cell/tissue types in mouse (A) or human (B) and tested in various cell/tissue types in human or mouse, respectively. The statistical tests were done using Wilcoxon signed-rank test. **C, D.** Boxplots of coefficients of the four marks in the model trained and tested in the same cell/tissue types in human (C) or mouse (D). **E, F.** Boxplots of coefficients of the four marks in the models trained in multiple cell/tissue types in human (E) or in mouse (F) and tested in different cell/tissue types in the same species. **p<0.001 (Wilcoxon signed-rank test), the weights of the labeled mark are significantly higher than those of the other three marks; ^ΔΔ^p<0.001 (Wilcoxon signed-rank test), the weights of the labeled mark are significantly higher than those of the other two marks. **G, H.** Coefficients of the four marks in the model trained in multiple cell/tissue types in mouse (G) or human (H) and tested in various cell/tissue types in human or mouse, respectively.

### Active proximal and distal CRMs can be largely differentiated by classifier models based on their epigenetic marks

We next asked whether the two categories of active CRMs can be discriminated based on their four epigenetic marks by a LR model trained on the active proximal CRMs as the positive sets and active distal CRMs as the negative sets. When trained and tested on the datasets from the same cell/tissue types, the models performed moderately well in differentiating active proximal CRMs and active distal CRMs in both human and mouse with a median AUROC of 0.798 and 0.815, respectively (Figures S12A, S12B). The models trained on n-1 cell/tissue types in the human and mouse also performed moderately well in the left-out cell/tissue types in the same species with a median AUROC of 0.787 and 0.788, respectively, which were significantly (p<1.41x10^-10^ and p<6.42x10^-11^) lower than those of the models trained and tested in the same cell/tissue types (Figures S12A, S12B). The model trained on mouse or human cell/tissue types also performed moderately well in human or mouse cell/tissue types with a median AUROC of 0.761 and 0.809, respectively, which were significantly lower (p<5.42x10^-11^, p<1.27x10^-7^) than those of the models trained and tested in cell/tissue types in the same species (Figures S12A, S12B). In almost all the models in the three scenarios, H3K4me3 had the highest positive weights (p<5.49x10^-9^) and H3K4me1 the lowest negative weights (p<0.001), while CA and H3K27ac had near zero weights (Figures S12C∼ S12H). These results suggest that the two categories of active CRMs have largely opposite patterns of H3K4me3 and H3K4me1 modifications, but largely indistinguishable CA and H3K27ac modifications, consistent with the current understanding of histone modifications on promoters and enhancers [54, 55, 67, 68]. Taken together, active proximal and distal CRMs might have distinct patterns of H3K4me1 and H3K4me3 modifications across various cell/tissue types in the same and even in different species, and they can be largely differentiated simply based on such differences by LR models, though the accuracy is not very high due probably to their often-overlapping functions [66, 72].

### Functional states in a cell/tissue type of all putative CRMs in the genome can be accurately predicted by a universal predictor using few epigenetic marks

As expected, the size of the CRM+TF^+^ set in a human (Figure 8A) or mouse (Figure S13A) cell/tissue type is highly variable, depending on the number of available TF-binding datasets in the cell/tissue types (Figures S1A, S1B). For example, dePCRM2 predicted the largest CRM+TF^+^ sets in the most well-studied cell/tissue types with the largest numbers of available TF ChIP-seq datasets, such as the K562, LNCaP and MCF5 human cell lines (Figures S1A, 8A) and mouse liver, macrophages and embryonic stem cells (ESC) (Figures S1B, S13A), while it predicted much smaller CRM+TF^+^ sets in most of the other cell/tissue types with fewer available TF ChIP-seq datasets (Figures S1, 8A, S13A). The results support our earlier argument that the functional states in most cell/tissue types of the most of the 1,225,115 and 798,258 putative CRMs in the human and mouse genomes, respectively, cannot be predicted by dePCRM2, thus, are unknown and needed to be predicted. Having demonstrated the power of our machine-learning classifier models to differentiate the positive and negative sets CRM+TF/Non-CRM or CRM+TF/CRM+S^-^ constructed in the 67 human and 64 mouse cell/tissue types with relatively larger numbers of available TF binding datasets, we asked whether the functional states in any cell/tissue type of all the putative CRMs in the genome, including those CRMs whose functional states cannot be predicted by dePCRM2 due to the lack of sufficient TF binding data [48], can be accurately predicted using data of a few epigenetic marks. To this end, since we have demonstrated in this study that at least human and mouse share the same epigenetic rules for defining functional states of putative CRMs in various cell/tissue types (Figures 5∼7), we constructed pairs of positive and negative sets by pooling all the positive sets CRM+TF^+^ and all the negative sets Non-CRM in the 67 human and 64 mouse cell/tissue types except the target cell/tissue type, respectively. Using these pairs of comprehensive positive and negative sets and the four epigenetic marks (CA, H3K4me1, H3K27ac and H3K4me3) as the features, we trained LR models called universal functional states predictors (UFSPs) of mammal CRMs. Using these UFSPs we predicted functional states in each of the 67 human and 64 mouse cell/tissue type of all the 1,225,115 and 798,258 putative CRMs in the human and mouse genomes, respectively (Methods).

**Figure 8.**
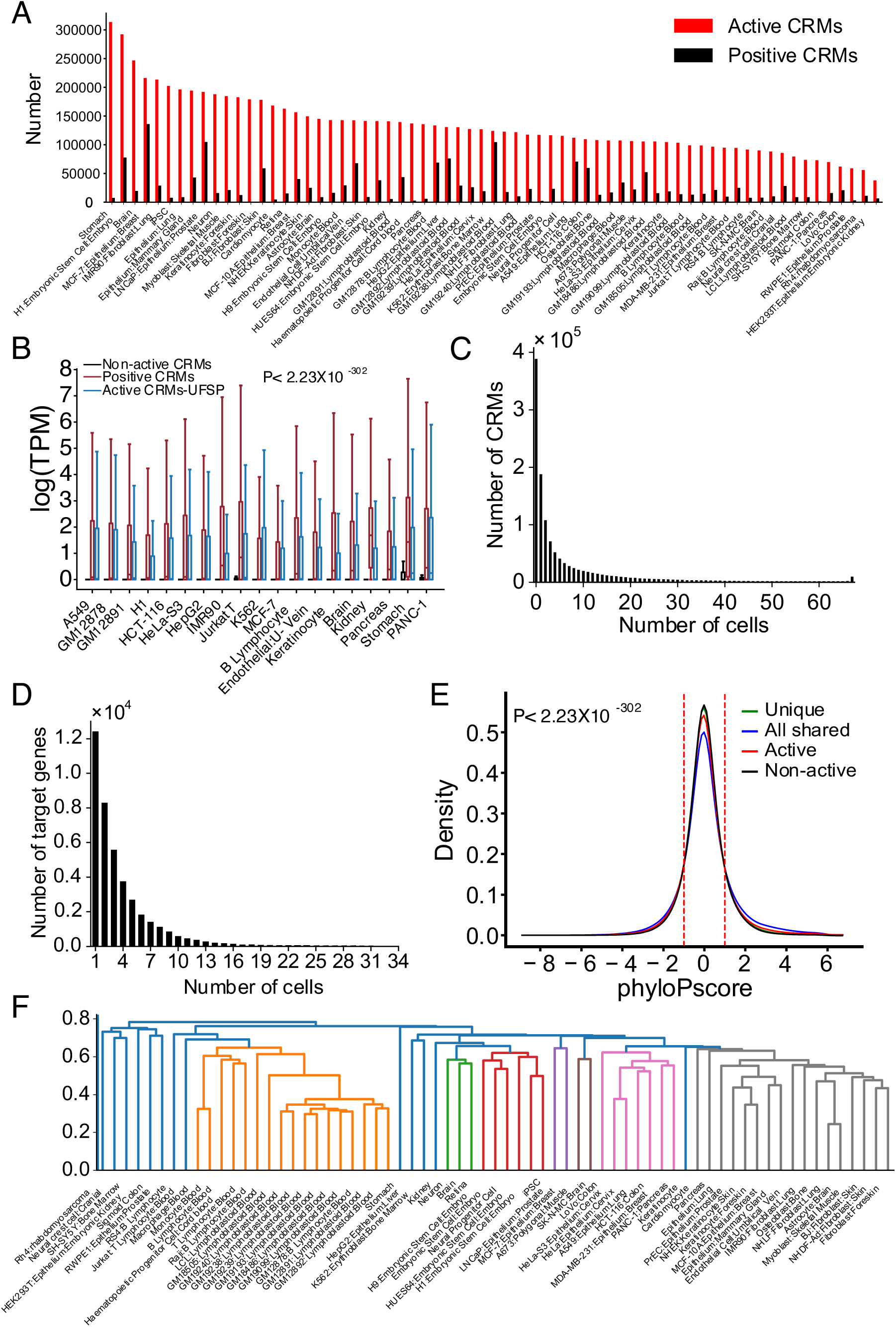
Genome-wide predictions of active CRMs in a human cell/tissue type and their reutilizations in different cell/tissue types. **A.** Number of active CRMs predicted by the UFSP (active CRMs) versus the size of the positive set CRM+TF+ (positive CRMs) in a cell/tissue type. **B.** Boxplots of gene expression levels in a cell/tissue type showing that genes closest to the CRM+TF^+^ set (positive) or to the active CRMs predicted by the UFSP model but missed by dePCRM2 (active CRMs-UFSP) have significantly higher expression levels than genes closest to the predicted non-active CRMs (p<2.23X10^-302^, Mann Whitney U test). **C.** Number of predicted active CRMs shared by different numbers of cell/tissue types. **D.** Number of closest genes to the uniquely active CRMs shared by different numbers of cell/tissue types. **E.** Distributions of phyloP scores of all-shared active CRMs, uniquely active CRMs, all active CRMs and all non-active CRMs in the cell/tissue types. All distributions are significantly different from one another, p<2.23X10^-302^ (K-S test). **F.** The levels of shared active CRMs reflect lineage relationships of the cell/tissue types. Cell/tissue types were clustered based on the Jaccard index of predicted active CRMs in each pair of the cell/tissue types.

The UFSPs predicted a highly varying number of active CRMs in a cell/tissue type, ranging from 37,792 (3.1%) to 313,389 (25.6%) and from 37,899 (4.8%) to 180,827 (22.7%), with a mean of 133,250 (10.9%) and 89,751 (11.2%) in the human and mouse cell/tissue types, respectively (Figures 8A, S13A). The UFSPs also predicted the remaining CRMs to be non-active in the cell/tissue types (Methods). As expected, in each cell/tissue type, the number of active CRMs predicted by the UFSP trained on all the other cell/tissue types is larger than the size of the positive set CRM+TF^+^ in the cell/tissue type (Figures 8A, S13A). Almost the entire positive set in the cell/tissue type, which was not used in training, was predicted to be active (data not show but see Figures 5A and 5B), suggesting that the UFSP is able to predict active CRMs in the cell/tissue type, which were missed by dePCRM2 using currently available TF-binding data. Notably, even in the most well-studied human cell lines such as K562, LNCaP and MCF5 (Figure S1) and mouse cell/tissue types such as live, macrophage and ESCs (Figure S2), UFSP was still able to predict active CRMs missed by dePCRM2 (Figures 8A, S13A). This result indicates that, even in these most ChIP-ed cell/tissue types, more TF-ChIP-seq datasets for more diverse TFs are needed to predict all active CRMs in them if only TF-ChIP-seq data are used, and thus, this method is high costly. Interestingly, in both human (Figure 8A) and mouse (Figure S13A), pluripotent embryonic stem cells and complex tissues (such as stomach and brain) generally had more active CRMs than purified terminally differentiated cell types (such as T or B lymphocytes). This is not surprising as pluripotent stem cells tend to express more genes than more differentiated cell types [73, 74], while the number of active CRMs in a complex tissue is the sum of active CRMs in each cell type in it that may contain many diverse cell types.

To verify the predictions, in each of the 19 human (Table S3) and 14 mouse (Table S4) cell/tissue types with RNA-seq data available, we split the active CRMs predicted by the UFSP into two sets: those that were also predicted to be active by dePCRM2 using available TF binding data (positive), and those that could not be predicted to be active by dePCRM2 due to the lack of sufficient TF binding data (active CRMs-UFSP). In each cell/tissue type, we then compared the expression levels of genes closest to the active CRMs-UFSP, positive CRMs (CRM-TF^+^) or non-active CRMs. We found that genes closest to the active CRMs-UFSP had similarly high expression levels as those closest to the positive CRMs, while both sets of genes had significantly higher (p<2.23X10^-302^) expression levels than those closest to the predicted non-active CRMs in all the 19 human (Figure 8B) and 14 mouse (Figure S13B) cell/tissue types. These results strongly suggest at least most of the active CRMs and non-active CRMs predicted by the UFSPs in both the human and mouse cell/tissue types might be authentic. Therefore, functional states in a cell/tissue type of all the putative CRMs in the genome can be accurately predicted by a universal classifier model using few epigenetic marks in the very cell/tissue type. The predicted results in the 67 human and 64 mouse cell/tissue types for all the 1,225,115 and 798,258 putative CRMs in the human and mouse genomes, respectively, are available at https://github.com/zhengchangsulab/Functional_states_of_CRMs.

### Most CRMs are extensively re-utilized in different cell/tissue types

Our predictions of the functional states of all the 1,225,115 and 798,258 putative CRMs in the human and the mouse genomes in the 67 human and 64 mouse cell/tissue types, respectively, positioned us to analyze the usage patterns of all the putative CRMs. We found that 68.28% (836,527) and 63.26% (505,016) of the putative CRMs in the human and mouse genome were predicted to be active in at least one of the 67 human (Figure 8C) and 64 mouse (Figure S13C) cell/tissue types, respectively. The remaining 31.72% and 36.14% of the putative CRMs were predicted to be non-active in all the human and mouse cell/tissue types analyzed, respectively. It is likely that these non-active CRMs are active in other cell/tissue types that were not analyzed in this study. Interestingly, of all the predicted active CRMs in the 67 human (n=836,527) and 64 mouse (n=505,016) cell/tissue types, only 22.44% (187,688) and 20.61% (104,074) were used in a single cell/tissue type, while the remaining 77.56% and 79.39% were reused in at least two cell/tissue types analyzed (Figure 8C, S13C), respectively, indicating that most CRMs were reused in the different human and mouse cell/tissue types. The number of uniquely active CRMs in a cell/tissue type ranged from 27 to 43,333 and from 32 to 6,914 with a mean of 2,801 and 1,626, comprising from 0.02% to 13.83% and from 0.08% to 5.62% of predicted active CRMs in a human (Figures S14A, S14B) and mouse (Figures S14C, S14D) cell/tissue type, respectively. Gene ontology (GO) term analysis [75–77] indicates that genes closest to uniquely active CRMs in a cell/tissue type are involved in functions specific to the cell/tissue type. For example, uniquely active CRMs in human H1-hESCs (H1) are closest to genes enriched for 236 GO terms for developments, such as tongue development (GO:0043586), establishment of epithelial cell polarity (GO:0090162), and negative regulation of axon extension (GO:0030517), to name a few (Table S5), while uniquely active CRMs in human brain tissues are closest to genes enriched for 177 GO terms for neuronal functions, such as inhibitory synapse assembly (GO:1904862), neuron cell-cell adhesion (GO:0007158), and ionotropic glutamate receptor signaling pathway (GO:0035235), etc, (Table S6). Similar results are seen for uniquely active CRMs in mouse embryonic stem cells (Table S7) and mouse brain cells (Table S8). These results suggest that a cell/tissue type might be determined by a set of the uniquely active CRMs in it. On the other hand, there were a total of 39,942 and 29,857 genes (including non-protein genes) closest to the uniquely active CRMs in the human and mouse cell/tissue types, however, only a total of 12,389 (31.0%) and 11,741 (39.2%) of them were unique to a human (Figure 8D) and mouse (Figure S13D) cell/tissue type, respectively. The remaining majority (69.0% and 60.8%) of genes were shared by at least two cell/tissue types, but no gene was shared by all the 67 human or 64 mouse cell/tissue types (Figures 8D, S13D). This result is in agreement with the notion that a cell type is determined by a unique combination of otherwise more widely expressed genes [78, 79].

Interestingly, in both the human and mouse cell/tissue types, the number of shared active CRMs decreased largely monotonously with the increase in the number of sharing cell/tissue types, with the exception that the number of active CRMs shared by all the cell/tissue types analyzed in human (9,537 (1.14%)) and in mouse (8,869 (1.11%)) was larger than that shared by some fewer numbers of cell/tissue types (Figures 8C, S13C). The 9,537 and 8,869 active CRMs shared by all the human and mouse cell/tissue types comprised from 3.04% to 25.25% and from 4.91% to 23.4% of active CRMs in a human and mouse cell/tissue type, respectively. GO term analysis [75–77] indicates that genes closest to these all-shared active CRMs are enriched for 929 and 859 GO terms for house-keeping functions in human (Table S9) and mouse (Table S10), respectively, such as amino acid activation (GO:0043038), cell death (GO:0008219), and ribosomal large subunit biogenesis (GO:0042273), to name a few, suggesting that the functions of all-shared CRMs are largely conserved.

The all-shared active CRMs are more likely under either positive selection or negative selection than the uniquely active CRMs (p<2.23X10^-302^) and the predicted non-active CRMs (p<2.23X10^-302^) as indicated by their respective phyloP score [64] distributions (Figured 8E and S13E). It is likely that these non-active CRMs might be uniquely active in or only shared by other cell/tissue types yet to be analyzed. To see to what extent the shared active CRMs in cell/tissue types reflect their developmental lineage relationships, we hierarchically clustered the cell/tissue types based on the Jaccard index of the predicted active CRMs of each pair of the cell/tissue types of an organism. As shown in Figures 8F and S13F, cell/tissue types with close lineages indeed formed clusters. These results support the notion that cell/tissue types are produced in a stepwise manner through cell differentiation, so that the closer cell types in a developmental lineage, the more gene regulatory programs they share [80–82].

### Our two-step approach is substantially more accurate and cost effective than state-of-the-art one-step approaches

Having demonstrated that the functional states in a cell/tissue type of all the putative CRMs in either human or mouse genome, predicted by dePCRM2 using more available TF ChIP-seq datasets can be very accurately predicted by the UFSP model using only four epigenetic marks, we finally compared the performance of our two-step approach with five state-of-the-art machine learning-based methods that attempt to predict active enhancers and/or promoters in a cell/tissue type using epigenetic marks in the very cell/tissue type (Methods). As summarized in Table 2, due to the lack of large sets of gold standard active CRMs and non-active CRMs in any animal cell/tissue types, to construct a positive set in a cell/tissue type, Matched Filter uses 2-kb genome regions overlapping STARR-seq and H3K27ac or CA peaks in the cell/tissue type, while the other earlier method generally use 2-kb regions overlapping histone acetyltransferase EP300 binding peaks in the cell/tissue type; to construct a negative set the cell/tissue type, all these earlier methods generally use randomly selected 2-kb bins without the features of the positive sets. Moreover, due to the lack of a map of CRMs in the genome, once trained, these earlier methods perform genome-wide predictions of active CRMs by evaluating a 2-kb sliding window, attempting to simultaneously predict the loci and functional states of CRMs in a given cell/tissue type (Table 2).

**Table 2.** Summary of methods for defining positive and negative sets for model training and candidate CRMs for genome-wide predictions.

| Methods | Labels | Positive set | Negative set | Classifier | Epigenetic marks data used | CRM candidates |
| --- | --- | --- | --- | --- | --- | --- |
| Our method | TF binding | CRMs overlapping TF-binding peaks | Randomly selected non-CRMs or CRM not overlapping STARR-seq peaks | LR | CA, H3K27ac, H3K4me1, H3K4me3 | Predicted CRMs |
| Matched Filter | STARR-seq & H3K27ac peaks | 2-kb regions around STARR-seq peaks overlapping H3K27ac or CA peaks | Randomly selected 2-kb bins not overlapping STARR and H3K27ac/CA peaks | SVM, random forest, rigid regression | CA, H3K9ac, H3K27ac, H3K4m1, H3K4m2, H3K4m3 | 2-kb sliding window |
| REPTILE | EP300 binding | DMRs in $\pm 1$-kb regions around the summits of top EP300 peaks | Randomly selected 2-kb bins not overlapping EP300 peaks | Random forest | mCG, H3K4me1, H3K4me2, H3K4me3, H3K27me3, H3K9ac, H3K27ac | 2-kb sliding windows with 100-bp step size |
| RF ECS | EP300 binding | $\pm 1$-kb regions around the summits of top EP300 peaks | Randomly selected 2-kb bins not overlapping EP300 peaks | Random forest | mCG, H3K4me1, H3K4me2, H3K4me3, H3K27me3, H3K9ac, H3K27ac | 2-kb sliding windows with 100-bp step size |
| DELTA | EP300 binding and promoters | Top EP300 peaks and all known promoter | Randomly selected 2-kb bins not overlapping EP300 peaks and promoters | AdaBoost | mCG, H3K4me1, H3K4me2, H3K4me3, H3K27me3, H3K9ac, H3K27ac | 2-kb sliding windows with 100-bp step size |
| CSI-ANN | EP300 binding or known enhancers | Known enhancers or top EP300 peaks | Randomly selected 2-kb bins | Neural network | mCG, H3K4me1, H3K4me2, H3K4me3, H3K27me3, H3K9ac, H3K27ac | 2-kb sliding windows with 100-bp step size |

We first compared the performance of our method with that of the five earlier methods on four mouse embryonic tissues including hindbrain, limb, midbrain and neural tube using training sets constructed (Table 2) and experimental data used in REPTILE [35] and Matched Filter [58]. As REPTILE [35], RFECS [41], DELTA [45] and CSI-ANN [40] models were trained on data from the mESCs [35], for a fairer comparison, we trained a RL model using the CRM+TF^+^/Non-CRM sets from the mESCs and pooled CRM+TF^+^/Non-CRM sets from the 67 human cell/tissue types. As shown in Figure 9A, both our mouse and human models achieved almost perfect AUROC values (0.992∼0.997) in the four mouse tissues, thus substantially outperforming all the five earlier classifiers (Figure 9B). This result is remarkable as we only used four epigenetic marks while all these earlier methods used more marks in addition to these four marks (Table 2). We attribute the superior performance of our method to the high accuracy of our predicted CRMs and non-CRMs, up on which high quality training sets can be possibly constructed. Notably, despite being trained on Drosophila S2 cells, Matched Filter outperformed the other four earlier methods on all the four tissues, supporting that the epigenetic rules defining functional states of CRMs might be universal from insects to mammals.

**Figure 9.**
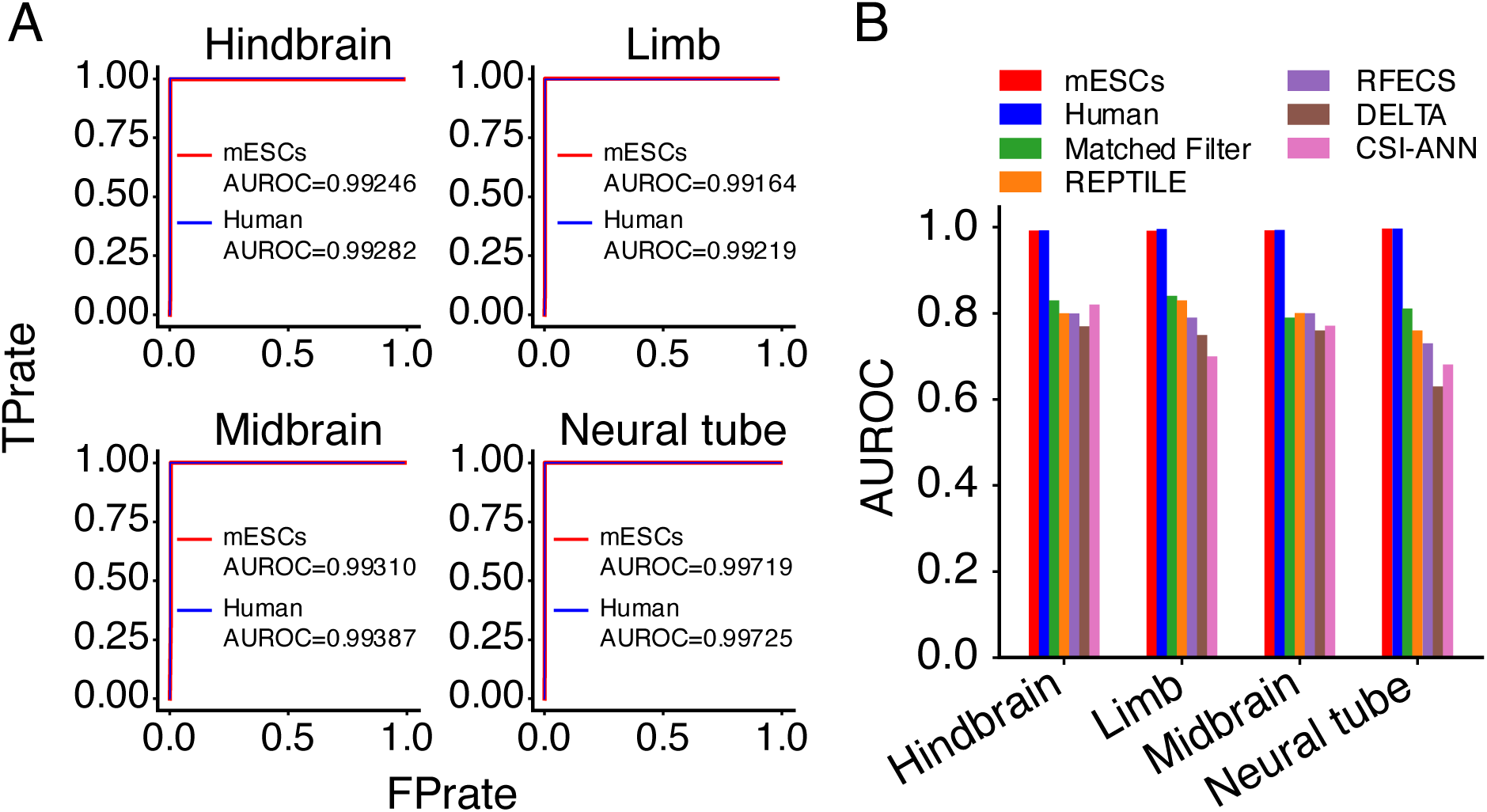
Comparison of the performance of our method with five earlier state-of-the-art methods on four mouse embryonic tissues. **A.** ROC curves in the mouse tissues of our LR model trained on the CRM+TF^+^/Non-CRM sets in the mESCs or in the 67 human cell/tissue types. Notably, the performance of the two models is almost indistinguishable. **B.** AUROCs in the mouse tissues of our LR models trained on the mESCs or on the human cell/tissue types in comparison with those of the five earlier methods. REPTIPLE, RFECS, DELTA and CSI-ANN models were trained on the positive/negative sets (Table 2) in the mESCs as reported in reference [35], and the Matched Filter model was trained on the positive/negativize sets (Table 2) in the *Drosophila* S2 cells as reported in reference [58]. Experimental data reported in references [35, 58] were used for the predictions in the four mouse tissues.

In the recent study [58], Matched Filter has been used for genome-wide predictions of active CRMs in six ENCODE top-tier human cell lines (H1-hESC, GM12878, K562, HepG2, A549 and MCF-7), we therefore also compared our predicted active CRMs in these cell lines using the LR model trained on pooled CRM+TF^+^/Non-CRM sets in the 64 mouse cell/tissue types. As shown in Figure 10A, our method predicted a much larger number of active CRMs in the six cell lines than did Matched Filter. Our predicted active CRMs in each cell line also cover a much larger proportion of the genome than did those predicted by Matched Filter (Figure 10B). Of the union of nucleotide positions covered by active CRMs in the six cell lines predicted by Matched Filter (213,326,167bp) and our method (873,473,948bp), only 82,414,070bp were predicted by both methods, comprising 38.63% and 10.42% of their predicted active CRM positions, respectively, while the remaining 61.37% and 89.58% were only predicted by Matched Filter and our method, respectively (Figure 10C). Thus, the vast majority of positions of active CRMs predicted by the two methods are different, although a small portion of them are the same. To see which method is more accurate than the other in predicting active CRMs in the six cell lines, we first analyzed phyloP conservation scores of positions shared by active CRMs predicted by both methods and of positions of active CRMs predicted only by one of the two methods. As shown in Figure 10D, positions of active CRMs predicted only by Matched Filter have a narrow, high peak distribution of their phyloP scores around 0, indicating that most of the positions are selectively neutral, and thus unlikely to be functional. In contrast, positions shared by active CRMs predicted by the two methods have a broad, low peak distribution of their phyloP scores around 0, and the same is true for positions of active CRMs predicted only by our method (Figure 10D), indicating that the vast majority of the both sets of positions are under evolutionary selections, and thus likely to be functional. Therefore, most positions (61.37%) of active CRMs predicted by Matched Filter are even not CRM loci at all. To further confirm this conclusion, we next compared the expression levels of genes closest to the active CRMs in a cell line predicted by each method, although the closest gene is not necessarily the target of an active CRM. As shown in Figure 10E, in all the six cell lines, genes closest to active CRMs predicted only by Matched Filter (no overlaps with any active CRMs predicted by our methods) had significantly lower expression levels than genes closest to the active CRMs predicted by both methods (the two putative active CRMs overlap at least half of the short one), as well as genes closest to the active CRMs only predicted by our method, suggesting that these active CRMs predicted only by Matched Filter might not be active or even not CRMs at all. Taken together, these results indicate that our two-step approach not only has achieved thus far the best performance for predicting both CRM loci in genomes and functional states of all the putative CRMs in a given cell/tissue type, but also is more cost effective than existing methods as it only needs four or even fewer epigenetic marks to achieve such high accuracy.

**Figure 10.**
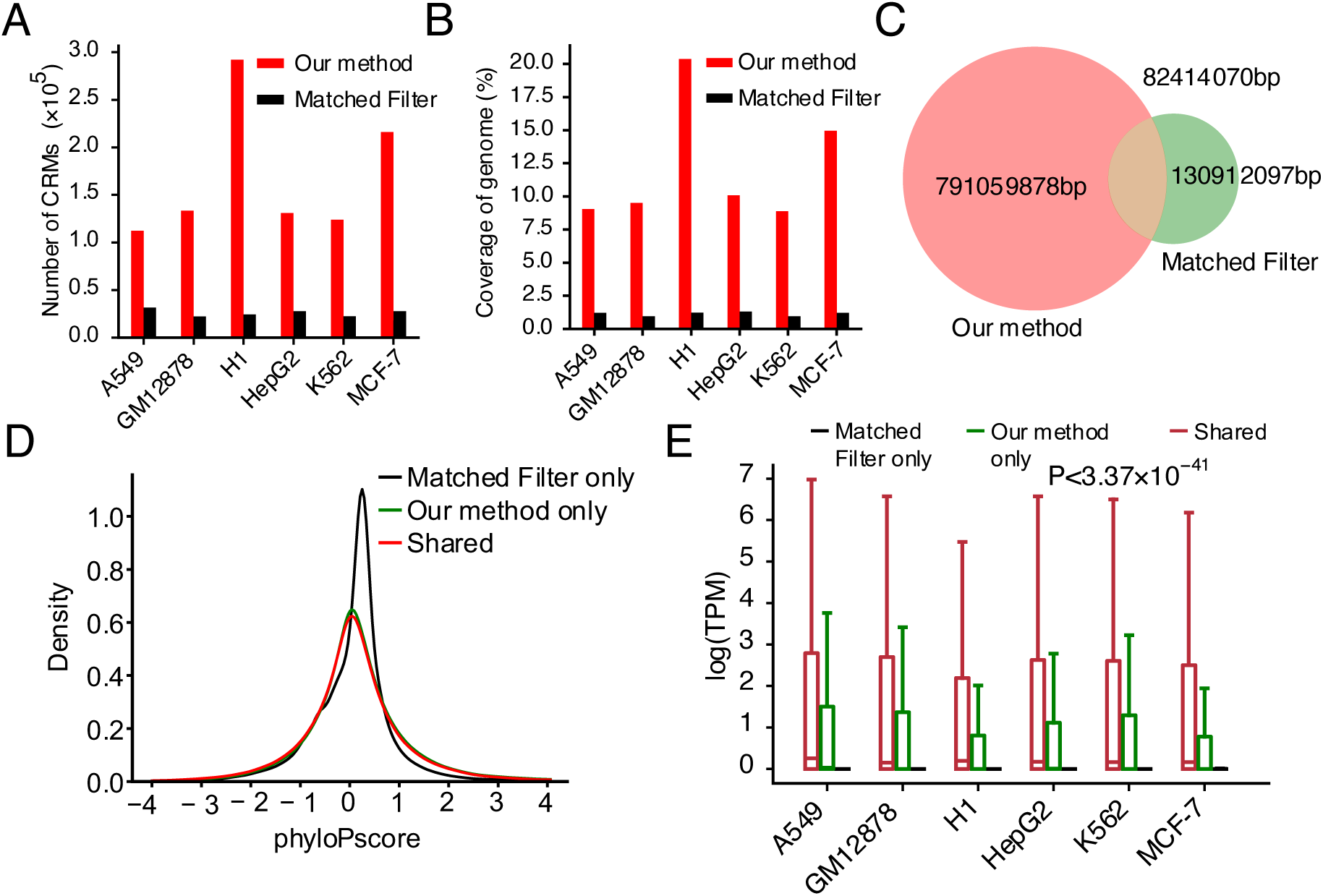
Comparison of the performance of our method and Matched Filter for genome-wide predictions of active CRMs in six human cell lines. **A.** Number of active CRMs predicted by the two methods in the cell lines. **B.** Proportion of the human genome covered by active CRMs predicted by the two methods in the cell lines. **C.** Venn diagram of the union of nucleotide positions covered by active CRMs predicted by each method in the six cell lines. **D.** Distributions of phyloP scores of nucleotide positions covered by active CRMs predicted by both methods (shared) and by active CRMs predicted only by Matched Filter (Matched Filter only) or only by our method (our method only). **E.** Boxplots of the expression levels of genes closest to active CRMs predicted by both methods (overlapping each other by at least 50% of the shorter one) in a cell line (shared) and of genes closest to active CRMs predicted only by Matched Filter (Matched Filter only) or only by our method (our method only). In each cell/tissue type, genes closest to active CRMs predicted by both methods and genes closest to active CRMs predicted only by our method have significantly higher expression levels than genes closest to active CRMs predicted only by Matched Filter (p<3.37X10^-41^, Mann Whitney U test).

## Discussion

Annotation of CRMs in a genome has three tasks. The first is to identify all CRMs and constituent TFBSs in the genome; the second is to characterize the functional state (active or non-active) of each CRM in each cell/tissue type of the organism; and the third is to determine the target genes of each active CRM in each cell/tissue type. The first and the second tasks are clearly two facets of the same coin, solving one would facilitate solving the other, and thus, it is attractive to solve them simultaneously. Indeed, most existing methods attempt to predict active CRMs in a cell/tissue type, and thus jointly predict CRM loci and their functional states in one step, by integrating epigenetic data in the very cell/tissue type using various machine-learning methods [36-43, 83-85]. Although conceptually attractive, these methods in practice have limitations due to the reasons we indicated earlier. In particular, a sequence segment with broader epigenetic marks such as H3K4me1 [52–54], H3K4me3 [55] and H3K27ac [56] or even narrow marks such as CA [15, 35] mGC [35], or their combinations [46-48, 56, 57] are not necessarily CRMs, although active CRMs do bear a certain pattern of them [26–31]. Moreover, it is difficult if not impossible, to de novo predict TFBSs in CRMs using histone marks and CA data alone. As a result, CRMs predicted by these methods are of low resolution with high FDRs [35, 43, 46–51] and lack information of constituent TFBSs, although some methods scan predicted CRMs for TFBSs of known motifs [85].

One way to circumvent the limitations of these methods might be to integrate epigenetic marks data with TF ChIP-seq data in a cell/tissue type, since it has been shown that an active CRM can be more accurately predicted using information of both chromatin modifications and bindings of key TFs [43, 46, 47, 51, 62]. However, the application of this approach is limited because sufficient TF ChIP-seq data are available only in very few well-studied cell lines [48]. A more cost-effective way to circumvent the limitations of these existing methods might be to take a two-step approach as we proposed earlier [48, 60] and fully implemented in this study (Figure 1). By developing the new pipeline dePCRM2 [48], we have provided a more efficient method for the first step to predict a highly accurate and more complete, yet largely cell/tissue type agnostic map of CRMs and constituent TFBSs in the genome at single-nucleotide resolution by integrating all available ChIP-seq datasets for different TFs in various cell/tissue types of the organism. We have shown that even using a relatively smaller number (6,097) of TF ChIP-seq datasets then available, the putative CRMs predicted by dePCRM2 in the human genome are more accurate and complete than those predicted by existing state-of-the-art methods such as SCREEN [86], EnhancerAtlas [84] and GeneHancer [85] that integrates enhancers predicted by ChromHMM [87] and Segway [38] in various cell/tissue types. In this study, we first predicted even more complete maps of putative CRMs in both the human and mouse genome using much larger numbers of TF binding datasets only available recently. We then presented a method for the second step of our approach to predict functional states in any cell/tissue type of all the putative CRMs in the genomes using optimal minimal sets of epigenomic marks data in the cell/tissue type. The rational of our method is the observation that once the locus of a CRM is accurately anchored by key binding TFs, its epigenetic marks in a cell/tissue type can be an accurate predictor of its functional state in the cell/tissue type [43, 46, 47, 62].

We showed that this two-step approach achieved substantially higher accuracy for predicting the functional states in both mouse and human cell/tissue types of all the putative CRMs than the existing state-of-the-art methods (Figures 2, 9,10). We attribute the outstanding performance of our methods to two novelties. First, based on the active CRMs predicted by dePCRM2 in a cell/tissue type with a relatively large number of TF ChIP-seq datasets available, we are able to construct a relatively large high-quality positive set CRM+TF^+^ and negative set CRM+S^-^ if STARR-seq data are available in the cell/tissue type; and if STARR-seq data are unavailable in the cell/tissue type, we can construct a large high quality negative set Non-CRM (Table 1). Importantly, we showed that these two negative sets (CRM+S^-^ or Non-CRM) have virtually indistinguishable patterns of light modifications of the four epigenetic marks (CA, H3K4me1, H3K4me3 and H3K27ac) analyzed, while the positive set CRM+TF^+^ show distinct patterns of heavy modifications of the four epigenetic marks from those of the two negative sets (Figures 3, S3, S5). By contrast, the positive and negative sets constructed by other methods (Tables 1) show less distinct patterns of the epigenetic marks (Figures 3, S3, S5). Therefore, it is not surprising that the LR models trained on the CRM+TF^+^/CRM+S^-^ sets or on the CRM+TF^+^/non-CRM sets performed equally well, and both models substantially outperformed those trained on the positive and negative sets constructed using other methods (Tables 1, and Figure 2A). It also is understandable that models of all the seven machine-learning classifiers trained on the positive sets CRM+TF^+^ and negative sets CRM+S^-^ (or non-CRM) performed almost equally very well (Figure 2E). It is also not surprising that our LR model trained on the CRM+TF^+^/Non-CRM sets in the mESCs or human cell/tissue types substantially outperformed on the four mouse embryonic tissues the earlier five machine-learning methods (Figure 9) trained on their positive and negative sets, which were typically constructed using 2-kb regions overlapping or not overlapping EP300 binding peaks or STARR-seq peaks (Table 2).

Second, dePCRM2 provides us a highly accurate and unprecedentedly complete maps of putative CRMs in 85.5% and 79.9% regions of the human and mouse genomes, respectively. Using these putative CRMs as candidate makes our genome-wide predictions of active CRMs in a cell/tissue less challenging, because it has been shown that once the locus of a CRM is anchored by the binding of key TFs, epigenetic marks on the CRM become an accurate predictor of its functional state [43, 46, 47, 51, 62]. By contrast, without such maps of putative CRMs, the existing methods generally use a 2-kb sliding window with a step size 100bp to scan the genome for predicting active CRMs in a cell/tissue type (Table 2), making the task more challenging. Indeed, our LR model that use the 1.2M putative CRMs predicted by dePCRM2 at p-value cutoff 0.05 in the human genome as CRM candidates, substantially outperformed the best earlier method Matched Filter that used 2-kb sliding windows as CRM candidates, for genome-wide predictions of active CRMs in the six human cell lines (Figure 10), although the different training sets (mouse data vs Drosophila data) used by the two methods might also contribute to the performance discrepancies.

Although dozens of epigenetic marks have been found to modify CRMs in different cellular contexts [58, 88], it remains elusive which and how many of them are required to define functional states of CRMs [35, 46, 54–56]. Recently, it was found that machine-learning models trained using as few as six marks (H3K27ac, H3K4me1, H3K4me2, H3K4me3, H3K9ac and CA) performed equally well as the models trained using as many as 30 marks in differentiating STARR-seq peaks and negative control sequences, with H3K27ac being the most important feature [58]. In this study, we show that functional states in a cell/tissue type of all the putative CRMs in the genome can be very accurately (AUROC>0.95) predicted using peaks of optimal minimal sets of one (CA), two (CA+H3K4me1), three (CA+H3K4me1+H3K4me3, or CA+H3K4me1+H3K27ac) and four (CA+H3K4me1+H3K4me3+H3K27ac) epigenetic marks with data widely available in cell/tissue types (Figures 4A, S8A). Using more than four epigenetic marks data could only infinitesimally increase the accuracy due to the redundant information in the data as indicated by the positive correlations between the peak signals of the six marks analyzed (Figures S7, S10). Therefore, once a map of CRMs in a genome is highly accurately and more completely predicted, our two-step approach can be highly cost-effective for predicting functional states in any cell/tissue type of all the putative CRMs in the genome by generating data of few (1∼4) epigenetic marks in the very cell/tissue type, although the more marks data used, the higher accuracy achieved. Our identified optimal minimal sets of marks might suggest a prioritization of data generation (Figures 4A, S7B, S8A, S10B).

Furthermore, we show that machine-learning models trained on pooled positive and pooled negative sets from multiple cell/tissue types in human or mouse can accurately predict functional states of CRMs in other cell/tissue types in the same species as well as in various cell/tissue types in the other species. These results confirm that the epigenetic rules for defining functional states of CRMs are common in developmentally closely related cell/tissue types in human and mouse [35] as well as from insects to mammals [58]. However, we found that the most critical epigenetic mark for defining functional states of CRMs is CA, rather than H3K27ac as suggested earlier [58]. On the contrary, our results show that H3K27ac is one of the three less important marks among the six marks analyzed (Figure S7B, S10B), consistent with a recent report that H3K27ac is dispensable in mouse embryonic stem cells [56]. It is highly likely that the earlier conclusion [58] was erroneously drawn since H3K27ac was used both as a feature and as the label in the training datasets as we replicated in this study (Figures 2A, 2C).

It is worth noting that the machine-learning classifiers actually differentiate the different labels on the positive and negative sets as defined in Tables 1 and 2. For instance, a classifier trained on the CRM+TF^+^/CRM+S^-^ sets (Table 1) in a cell type differentiated CRMs with TF binding from CRMs without STARR-seq signals. In other words, the classifier was trained to predict whether a CRM was bound by TFs or did not overlap a STARR-seq peak in the cell/tissue type, given the epigenetic profile of the CRM. Thus, the labels may not necessarily reflect the activities of the CRMs. However, before the availability of large gold standard active and non-active CRMs sets in a cell/tissue type, such operational definition of positive and negative sets using a certain label on candidate CRMs might be the only choice that one can use to train machine-learning models. Nonetheless, although TFs binding to a silencer might reduce gene expression, while TFs binding to an enhancer may not necessarily enhance gene expression, we found that genes closest to the putative CRMs in the positive sets CRM+TF^+^ (Figure 2B) and the predicted active CRMs (Figure 8B, S13B) have significantly higher expression levels than those closest to the sequences in the negative sets non-CRM or CRM+S^-^ (Figure 2B) and the predicted non-active CRMs (Figure 8B, S13B), which had very low expression levels. These results strongly suggest that putative CRMs in the positive sets CRM+TF^+^ and the predicted active CRMs tend to enhance gene expression, while sequences in the negative sets and predicted non-active CRMs tend to not. Therefore, our definitions of active and non-active states largely reflect the functional states of CRMs.

The high accuracy of our predicted active CRMs positioned us to address two related interesting questions. 1) How many of the 1.2M and 0.8M CRMs predicted by dePCRM2 in the human and mouse genomes, respectively, are active in a cell/tissue type of the organisms, and 2) how many active CRMs are needed to define a cell/tissue type? We found that different cell/tissue types of humans and mice have widely varying numbers of active CRMs, ranging from 37,792 to 313,389 and from 37,899 to 180,827, respectively, depending on their cellular complexity and differentiation stages. Of these active CRMs, from only 27 (0.02%) to 43,333 (13.83%) and from only 32 (0.08%) to 6,914 (5.62%) are unique to a human and mouse cell/tissue type, respectively. We show that genes closest to the uniquely active CRMs are enriched for GO terms related to the functions of the cell/tissue. Thus, it appears that uniquely active CRMs in a cell/tissue type largely specify the cell/tissue type.

Moreover, only a third of genes closest to uniquely active CRMs are unique to a cell/tissue type (Figures 8D, S13D), supporting the notion that a terminally differentiated cell type is determined by a unique combination of otherwise more widely expressed genes [78, 79]. In this regarding, we note that the human genome encodes 33.4 times CRMs (n=1.47M) [48] as genes (n=44K), making it possible to use a set of uniquely active CRMs to regulate a specific combination of otherwise more widely expressed genes that ultimately determine the cell type. On the other hand, the vast majority of active CRMs in a cell/tissue type are reutilized in at least one of the other cell/tissue types analyzed. However, since many complex tissues in our analysis, such as brain, testis and stomach, to name a few, might contain multiple cell types, and since the numbers of cell/tissue types we analyzed are still small, it is likely that we might have overestimated the upper bounds of uniquely active CRMs in both the human and mouse cell types. Interestingly, cell/tissue types with related lineages form clusters based on the extent to which they share active CRMs (Jaccard index). This result is consistent with the notion that cell types are produced in a stepwise manner during cell differentiation, and thus, cell types that differentiate more recently from the last common ancestral type share more active CRMs [80–82].

### Conclusions

We present a two-step approach to predict functional states in any cell/tissue type of all putative CRMs using optimal minimal sets of four widely available epigenetic marks in the very cell/tissue type. Our approach substantially outperforms existing state-of-the-art methods that attempt to jointly predict CRM loci and their functional states in a given cell type using at least six epigenetic marks. The next step would be to develop a method to more accurately predict the target genes of active CRMs in a cell/tissue type beyond the “closest gene principle” as used in this study and others [89].

## Methods

### The datasets

We downloaded from CISTROME (http://cistrome.org/db/<u>#/</u>) [90] (12/20/2020) 11,348 and 9,060 TF ChIP-seq binding peak files for 1,360 and 701 TFs in 722 and 569 human (Table S1) and mouse (Table S2) cell/tissue types, respectively. Of these human and mouse cell/tissue types, 67 (Table S3) and 64 (Table S4), respectively, are most well-studied with ChIP-seq datasets available for four epigenetic marks (CA (measured by DNase-seq or ATAC-seq), H3K4me1, H3K27ac and H3K4me3) and varying number of TFs (Figure S1). Of the 67 human and 64 mouse cell/tissue types, 22 (Table S3) and 29 (Table S4), respectively, also have data available for two additional epigenetic marks (H3K4me2 and H3K9ac), and 19 (Table S3) and 14 (Table S4), respectively, have RNA-seq datasets available. We downloaded from CISTROME peak files of these epigenetic marks in the cell/tissue types. All the TF binding, CA and histone mark peaks were uniformly produced by the CISTROME team using the peak-calling tool MACS [91]. Of the 67 human cell/tissue types, six cell lines (A549, HCT116, HepG2, K562, MCH-7 and SH-HY5Y) have WHG-STARR-seq data available (Table S3). We downloaded gene expression data and WHG-STARR-seq peaks from the ENCODE data portal (https://www.encodeproject.org/). We downloaded experimental data in mouse embryonic tissues using access numbers and website links provided in Tables S3 and S4 in reference [35]. We downloaded predicted active enhancers and promoters in six human cell lines (H1-hESC, GM12878, K562, HepG2, A549 and MCF-7) [58] from https://github.com/gersteinlab/MatchedFilter.

### *De novo* genome-wide prediction of CRMs in the human and mouse genomes

For each called binding peak in each TF ChIP-seq dataset, we extracted 1,000bp genomic sequence centering on the summit of the peak. As most of the called peaks were shorter than 500bp (data not shown, but see [48]), we extended most of the called binding peaks. We have shown earlier that such extension could greatly increase the power of the datasets without including much noise [48, 92] (also see RESULTS). We applied dePCRM2 [48] to the extended binding peaks in the 11,348 and 9,060 datasets in the human and mouse cell/tissue types to predict the loci of CRMs and non-CRMs in the human and mouse genome regions covered by the extended binding peaks, respectively. Using DePCRM2, we also predicted a CRM to be active (TF-binding) in a cell/tissue type if at least one of constituent TFBSs in the CRM overlap the summit of a binding peak of a ChIP-ed TF in the cell/tissue type [48].

### Construction of a positive and negative CRM set in a cell/tissue type

We evaluated the following seven methods for constructing a positive (active) set and a negative (non-active) set of CRMs in a cell/tissue type (Table 1).

1) CRM+TF^+^/Non-CRM, where the positive set CRM+TF^+^ in a cell/tissue type consists of the active CRMs predicted by dePCRM2 in the cell/tissue type, and the negative set Non-CRM is randomly selected putative non-CRMs in the genome, with the matched number and lengths of sequences in the positive set (Table 1). The CRM+TF^+^ set is generally a small portion of all active CRMs in the cell/tissue type (see later). Clearly, we should not consider to be non-active a CRM that does not overlap any available TF binding peak in the cell/tissue, because the CRM may be bound by a TF that has not been ChIP-ed, and thus is actually active.
2) CRM+TF^+^/CRM+S^-^, where the positive set CRM+TF^+^ in a cell/tissue type is defined in the same way as above, and the negative set CRM+S^-^ is randomly selected putative CRMs in the genome that are not predicted to be active by dePCRM2 and does not overlap any STARR-seq peaks in the cell/tissue type, with the matched number and lengths of sequences in the positive set.
3) CRM+S^+^/Non-CRM, where the positive set CRM+S^+^ in a cell/tissue type consists of our predicted CRMs in the genome that overlap STARR-seq peaks in the cell/tissue type, and the negative set Non-CRM is randomly selected from the putative non-CRMs in the genome, with the matched number and lengths of sequences in the positive set.
4) CRM+S^+^/CRM+S^-^, where the positive and negative sets in a cell/tissue type are constructed in the same ways as in 3) and 2), respectively, with the negative set having the matched number and lengths of sequences in the positive set.
5) Bin+S^+^/Bin+S^-^, where the positive set Bin+S^+^ in a cell/tissue type is formed by 700bp genomic sequence bins that overlap STARR-seq peaks in the cell/tissue type, and the negative set Bin+S^-^ is randomly selected 700bp genomic bins that did not overlap any STARR-seq peaks in the cell/tissue type, with the matched number and lengths of sequences in the positive set.
6) Bin+ac^+^/Bin+ac^-^, where the positive set Bin+ac^+^ in a cell/tissue type is formed by 700bp genomic bins that overlap H3K27ac peaks in the cell line, and the negative set Bin+ac^-^ is randomly selected 700bp genomic bins that does not overlap any H3K27ac peaks in the cell/tissue type, with the matched number and lengths of sequences in the positive set. A similar method was used in an earlier study [59].
7) Bin+S^+^&ac^+^/Bin+S^-^&ac^-^, where the positive set Bin+S^+^&ac^+^ in a cell/tissue type is formed by 700bp genomic bins that overlap both H3K27ac and STARR-seq peaks in the cell line, and the negative set Bin+S^-^&ac^-^ is randomly selected 700bp genomic bins that overlap neither H3K27ac nor STARR-seq peaks in the cell/tissue type, with the matched number and lengths of sequences in the positive set. A similar method was used in an earlier study [58]. As the number and lengths of sequences in a negative set match those of the cognate positive set, all pairs of the positive and negative sets are well-balanced.

### Computing the epigenetic feature vector of a sequence

Given a sequence *t* (a predicted CRM, non-CRM or a genomic sequence bin), we compute a *M*-element raw feature vector *S_raw_*(*t*) for *M* epigenetic marks. If *t* overlaps at least one peak of the *m*-th epigenetic mark by at least 50% of the length of the shorter one, then the *m*-th element of *S_raw_*(*t*) is defined as

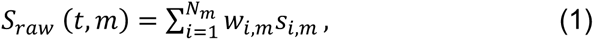

where *N_m_* is the number of peaks of the *m*-th mark overlapping *t* by at least 50% of the length of the shorter one, *w_i,m_* the ratio of the length of *t* over the length of the *i*-th peak of the *m*-th mark, *s_i,m_* the MACS signal score [91] of the *i*-th peak of the *m*-th mark. Clearly, if *t* does not overlap any peak of the *m*-th mark by at least 50% of the length of the shorter one, *S_raw_*(*t*, *m*) = 0. We then normalize the values of each *m*-th mark by

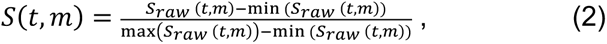

where min(*S_raw_*(*t*, *m*)) and max(*S_raw_*(*t*, *m*)) are the minimum and maximum of *S_raw_* (*t*, *m*) over all *t*, respectively. We use the normalized feature vectors *S*(*t*) to train machine-learning models and predict functional (TF binding) states of sequences.

### Model training, testing and evaluation

We evaluated seven machine-learning classifiers, including logistic regression, AdaBoost, SVM, neural network, naïve Bayes, decision tree and random forest. In a cell/tissue type, we trained a classifier model on a pair of positive and negative sets defined in the cell/tissue type (Table 1) using their normalized feature vectors of epigenetic marks. Shown in Figure 1 is the workflow of our machine-learning classifier models trained on the CRM+TF^+^/Non-CRM sets or on the CRM+TF^+^/CRM+S^-^ sets of a cell/tissue type. We conducted 10-fold cross-validation of the model in the cell/tissue type. In addition, in each species (human or mouse), we constructed a positive set and a negative set by pooling positive sets and negative sets in all cell/tissue types in the species, respectively. We trained a classifier model on the pooled human or mouse positive and negative sets using epigenetic marks as the features. We conducted leave-one-out cross-validation of a model in a species (human or mouse) by training the model on datasets in *n* − 1 cell/tissue types, and testing the model on the left-out cell/tissue type. We assessed the performance of a model using the area under the ROC (receiver operator characteristic) curve (AUROC), because each pair of the positive set and the negative set were well-balanced in both the number and lengths of sequences. We also evaluated the importance of epigenetic marks using their coefficients in logistic regression and SVM models. All of the classifier models were implemented using sci-kit learn v.0.24.2, and the code is available at https://github.com/zhengchangsulab/Functional_states_of_CRMs.

### Prediction of functional states in a cell/tissue type of all the CRMs in the genome

To predict the functional states in a given human or mouse target cell/tissue type of all the putative CRMs in the human or mouse genomes, we trained a LR model on the pooled positive (CRM+TF^+^) and pooled negative (Non-CRM) sets from the 67 human and 64 mouse cell/tissue types, except the target cell/tissue type using four epigenetic marks (CA, H3K4me1, H3K27ac and H3K4me3). We call this model a universal functional states predictor (UFSP) of mammal CRMs. Given a cell/tissue type with data for the four epigenetic marks, we applied UFSP to each of the putative CRMs in the genome, and predicted a CRM to be active (TF-binding) in the cell/tissue type if the CRM’s LR value ≥ 0.5, or non-active (non-TF-binding), otherwise.

### Comparison with other state-of-the-art methods

We compared our two-step approach with six earlier machine-learning methods that aim to simultaneously predict the loci and functional states of CRMs in a given cell/tissue type using epigenetic data from the very cell/tissue types. These methods include Matched Filter, a most recent method that combines a signal processing technique called matched filter [93] with a linear SVM, random forest or ridge regression model [58]; REPTILE [35], recent random forest based models that integrate histone modifications and bisulfite sequencing data for mGC modifications; RFECS [41], an earlier random forest based model; DELTA, an AdaBoost based ensemble method [45]; and CSI-ANN, a neural network based method [40]. For a fair comparison, we evaluated the performance of our methods and these earlier methods on the four mouse embryonic tissues (neural tube, midbrain, hindbrain and limb) and/or six ENCODE top-tier human cell lines (H1-hESC, GM12878, K562, HepG2, A549 and MCF-7) using training sets constructed and experimental data used in REPTILE [35] and Matched Filter [58].

### Declarations

**Ethics approval and consent to participate**

Not applicable

### Consent for publication

Not applicable

### Availability of data and materials

The software and predicted active CRMs in human and mouse cell/tissue types from the current study are available at https://github.com/zhengchangsulab/Functional_states_of_CRMs

The newly predicted CRMs in the human and mouse genomes in the current study are available from the corresponding author on reasonable request.

The TF ChIP-seq binding peaks and epigenetic mark peaks data used in the current study are available at http://cistrome.org/db/#/

The predicted active enhancers and promoters in six human cell lines (H1-hESC, GM12878, K562, HepG2, A549 and MCF-7) used in the current study are available at https://github.com/gersteinlab/MatchedFilter.

The gene expression data and WHG-STARR-seq peaks used in the current study are available at https://www.encodeproject.org/

The epigenetic marks data in the four mouse embryonic tissues used in the current study are available at the website links provided in Tables S3 and S4 in reference [35].

### Funding

The work was supported by the US National Science Foundation (DBI-1661332). The funding bodies played no role in the design of the study and collection, analysis, and interpretation of data and in writing the manuscript.

### Authors’ contributions

ZS and PN conceived the project. ZS and PN developed the algorithms, PN carried out all computational experiments and analysis and JM preprocessed the datasets. PN and ZS wrote the manuscripts. All authors read and approved the final manuscript.

## Abbreviations

AUROC: area under receiver operator characteristic curve
ATAC: assay for transposase-accessible chromatin
ATAC-seq: assay for transposase accessible chromatin using sequencing
CA: chromatin accessibility
ChIP-seq: chromatin immunoprecipitation sequencing
CRM: cis-regulatory module
DNase-seq: DNase I hypersensitive sites sequencing
ESC: embryonic stem cells
mESC: mouse ESC
FDRs: false discovery rates
LR: logistic regression
mCG: cytosine methylation in CpG dinucleotide
MPRA: massively parallel reporter assays
ROC: receiver operator characteristic curve
SVM: support vector machine
TF: transcription factor
TFBS: TF binding site
STARR-seq: self-transcribing assay of regulatory regions sequencing
UFSPs: universal functional states predictors
WHG- STARR-seq: whole genome STARR-seq.

## Acknowledgements

We would like to thank all lab members for their discussion, particularly, Sisi Yuan for her critical reading and suggestions.

## Supplementary Data

Supplementary tables S1∼S10 are available in Additional File 1, and supplemental figures S1∼14 are available in Additional File 2.

